# Benzylic Trifluoromethyl Accelerates 1,6-Elimination Toward Rapid Probe Activation

**DOI:** 10.1101/2024.05.30.596105

**Authors:** Liming Wang, Aditya Sivakumar, Rui Zhang, Sarah Cho, Yuhyun Kim, Tushar Aggarwal, Lu Wang, Enver Cagri Izgu

**Author notes:** Corresponding author: Izgu, E. C. These authors contributed equally (Rui Zhang, Sarah Cho).

## Abstract

Activity-based detection of hydrogen sulfide in live cells can expand our understanding of its reactivity and complex physiological effects. We have discovered a highly efficient method for fluorescent probe activation, which is driven by H_2_S-triggered 1,6-elimination of an α-CF_3_-benzyl to release resorufin. In detecting intracellular H_2_S, 4-azido-(α-CF_3_)-benzyl resorufin offers significantly faster signal generation and improved sensitivity compared to 4-azidobenzyl resorufin. Computed free energy profiles for the 1,6-elimination process support the hypothesis that a benzylic CF_3_ group can reduce the activation energy barrier toward probe activation. This novel probe design allows for near-real-time detection of H_2_S in HeLa cells under stimulation conditions.

## INTRODUCTION

Hydrogen sulfide (H_2_S) has emerged as a key reactive sulfur species (RSS) that serves as a gasotransmitter in mammals^1^ and is involved in signaling pathways concerning neurological^2–5^ and cardiovascular^6,7^ processes. H_2_S is also a vasorelaxant (maintained at concentrations around 40 μM in mice serum^8^) and a neuroprotectant in Alzheimer’s disease.^9,10^ Importantly, however, overproduction of H_2_S has been associated with various diseases, including Down syndrome,^11,12^ type 1 and type 2 diabetes,^13^ and cancer.^14^ These underscore the delicate and critical nature of regulating intracellular H_2_S levels.

The multifaceted roles of H_2_S in biological processes has attracted increasing attention, especially in the context of activity-based sensing.^15–22^ As the emerging studies continue to characterize further roles of RSS in complex biological systems,^23,24^ it has become crucial to gain specific understanding of the production of H_2_S. There are four main enzymatic pathways that produce majority of H_2_S in mammalian cells: through the catalysis of cystathionine-γ-lyase (CSE), cystathionine-β-synthase (CBS), 2-mercaptopyruvate sulfurtransferase (3-MST), or cysteinyl-tRNA-synthetases (CARS).^25^ Both CSE and CBS use L-cysteine (L-Cys) as a substrate to catalyze reactions, which generate H_2_S as a byproduct.^26^ On the other hand, literature provides mixed reports on intracellular H_2_S concentration, which indicates that measuring it accurately has been a challenge. Deeper insights into the biological consequences of H_2_S and related sulfur species are needed at both fundamental and translational levels.^27,28^ Real-time detection of H_2_S can help further characterize of its reactivity and effects targeting distinct areas such as the pulmonary, cardiac, nervous, and hepatic systems.^29–31^. However, it remains a challenge to detect intracellular H_2_S with high sensitivity and at high temporal resolution.^32^

Herein, we describe a new approach for designing probes for specific and rapid detection of intracellular H_2_S (Figure 1). This method uses H_2_S-triggered 1,6-elimination of α-CF_3_-benzyl caging group. We experimentally and computationally show that the benzylic CF_3_ group outperforms its protio analog as a caging group for resorufin. Live HeLa cell imaging suggested that 4-azido-(α-CF_3_)-benzyl resorufin can make it possible to detect the prevalence of H_2_S in near-real-time.

**Figure 1.**
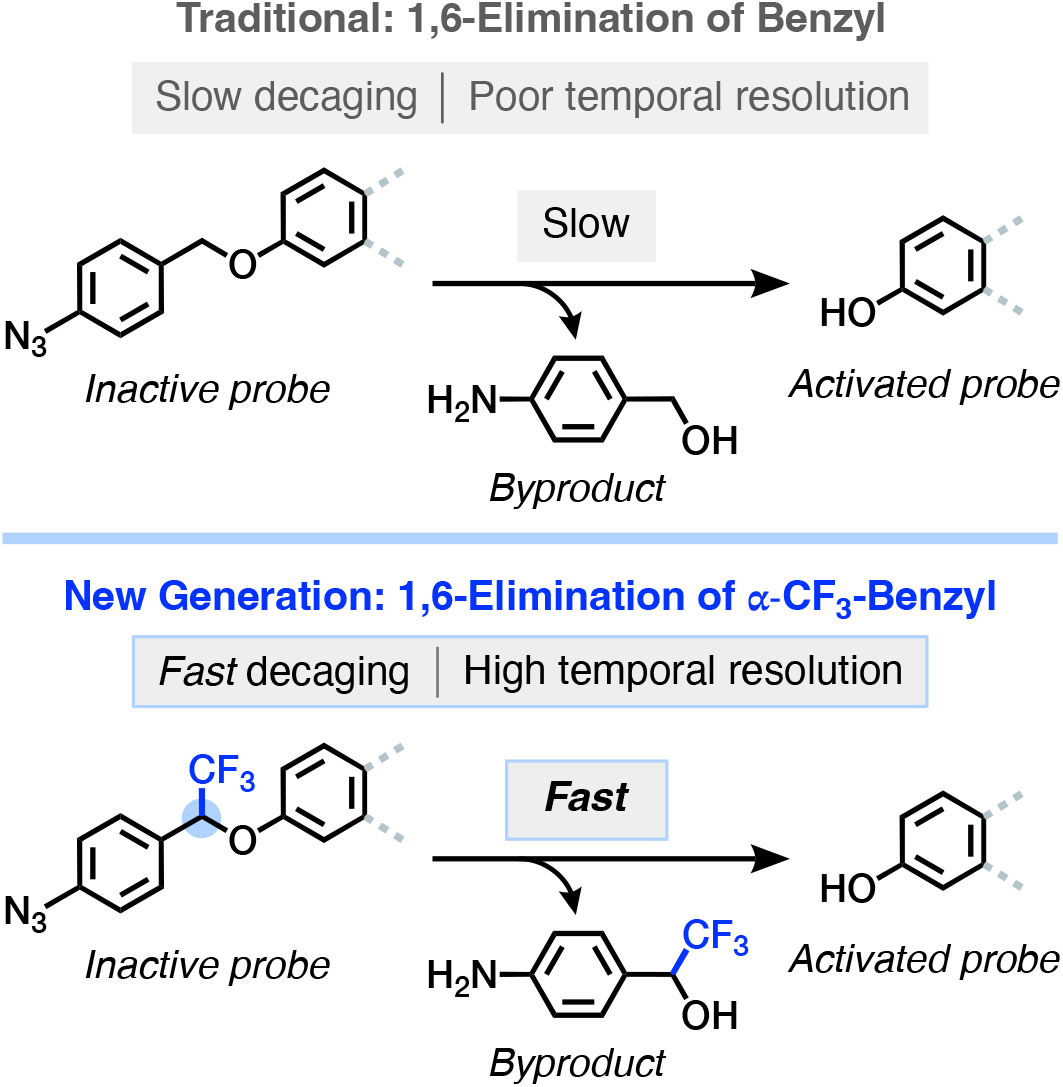
The concept of α-trifluoromethylation in 4-azidobenzyl-caged probe design. Traditional 4-azidobenzyl group undergoes a slow 1,6-elimination process (top), whereas the 4-azido-(α-CF_3_)-benzyl delivers a significantly faster 1,6-elimination, allowing for rapid probe activation (bottom).

## RESULTS AND DISCUSSION

### Designs and syntheses of resorufin-based H_2_S probes

In designing high-performance H_2_S probes, we reasoned that decreasing the activation energy barrier of 1,6-elimination can accelerate the fluorophore uncaging process. To this end, a viable approach would be to employ appendant electron-withdrawing group(s) to stabilize the transition state through which 1,6 elimination proceeds following azide reduction. Naturally, our preliminary designs included the substitution of benzylic protons for fluorine atoms, an isosteric electron-withdrawing group. This type of chemical transformation can be, in principle, delivered through established benzylic difluorination and substitution chemistry.^33^ However, a recent study on activity-based protein labeling employed quinone methide intermediates generated upon 1,6-elimination of benzylic fluoride.^34^ This brings forth α-F-benzylation in probe engineering as an unproductive strategy in terms of probe activation, necessitating the discovery of alternate elimination pathways. Our pursuit of finding this alternative elimination mechanism leveraged thermodynamic analyses of potential leaving groups at the benzylic (α) position, some considered atypical in organic synthesis. The p*K*_a_ of HF in water is 3,^35^ whereas the p*K*_a_ of resorufin in water is 5.8.^36^ This near 3 p*K*_a_ units of difference in acid strengths lends credence to the hypothesis that α-F-benzyl group would unlikely serve to uncage resorufin. In comparison, the p*K*_a_ of CF_3_H is 27,^37^ indicating that it is a far weaker acid than resorufin, and that its conjugate base, CF_3_^−^, is a substantially poorer leaving group. In addition, a comparison of the Hammett constants between the trifluoromethyl group (σ_p_ = 0.54) and fluorine (σ_p_ = 0.06) suggests the former has a stronger electron-withdrawing effect. These properties make CF_3_ group a promising substitution candidate at the benzylic position for 1,6-elimination reactions.^38^

The 7-*O*-caged resorufins ***7*** and ***8*** were obtained from resorufin (***6***) and the respective benzyl bromide, ***5*** or ***4*** (Figure 2). While 4-azidobenzyl bromide (***5***) is readily available, the α-CF_3_ analog ***4*** was synthesized in three linear steps from 4-azidobenzyl alcohol (***1***). Oxidation of ***1*** with Dess–Martin periodinane (DMP) provided 4-azidobenzaldehyde (***2***) in 82% isolated yield. This was followed by the functionalization of ***2*** with a CF_3_ group at the benzylic position. Using TMSCF_3_ as the source of CF_3_ and TBAF to breakdown the O-TMS ether generated upon trifluoromethylation, the alcohol ***3*** was formed in nearly quantitative yield (95%). Notably, the alcohol ***3*** was found to exhibit poor reactivity toward direct etherification. Extensive method exploration to determine a feasible strategy for the 4-azido-(α-CF_3_)-benzylation of resorufin (***6***) suggested nucleophilic substitution to be workable. In accordance, we converted the alcohol ***3*** to its brominated analog 1-azido-4-(1-bromo-2,2,2-trifluoroethyl)benzene (***4***) using P(OPh)_3_ and NBS in 81% yield. Subsequent 4-azidobenzylation of ***6*** with ***4*** in the presence of Cs_2_CO_3_ afforded the 4-azido-(α-CF_3_)-benzyl resorufin (***8***) in 10% yield. Though we can prepare ***8*** in quantities that enable downstream applications, the isolated product yield in this transformation is far from desired. Of note, this compound quickly degrades on silica gel column. We are concurrently exploring ways to increase the α-CF_3_-benzylation efficiency and will report our findings in due course. Finally, benzylation of ***6*** with 4-azidobenzyl bromide (***5***) delivered the 4-azidobenzyl resorufin (***7***) in 11% yield. This compound, like ***8***, was challenging to isolate through silica gel column chromatography. Therefore, its purification was carried out by trituration with ethyl acetate.

**Figure 2.**
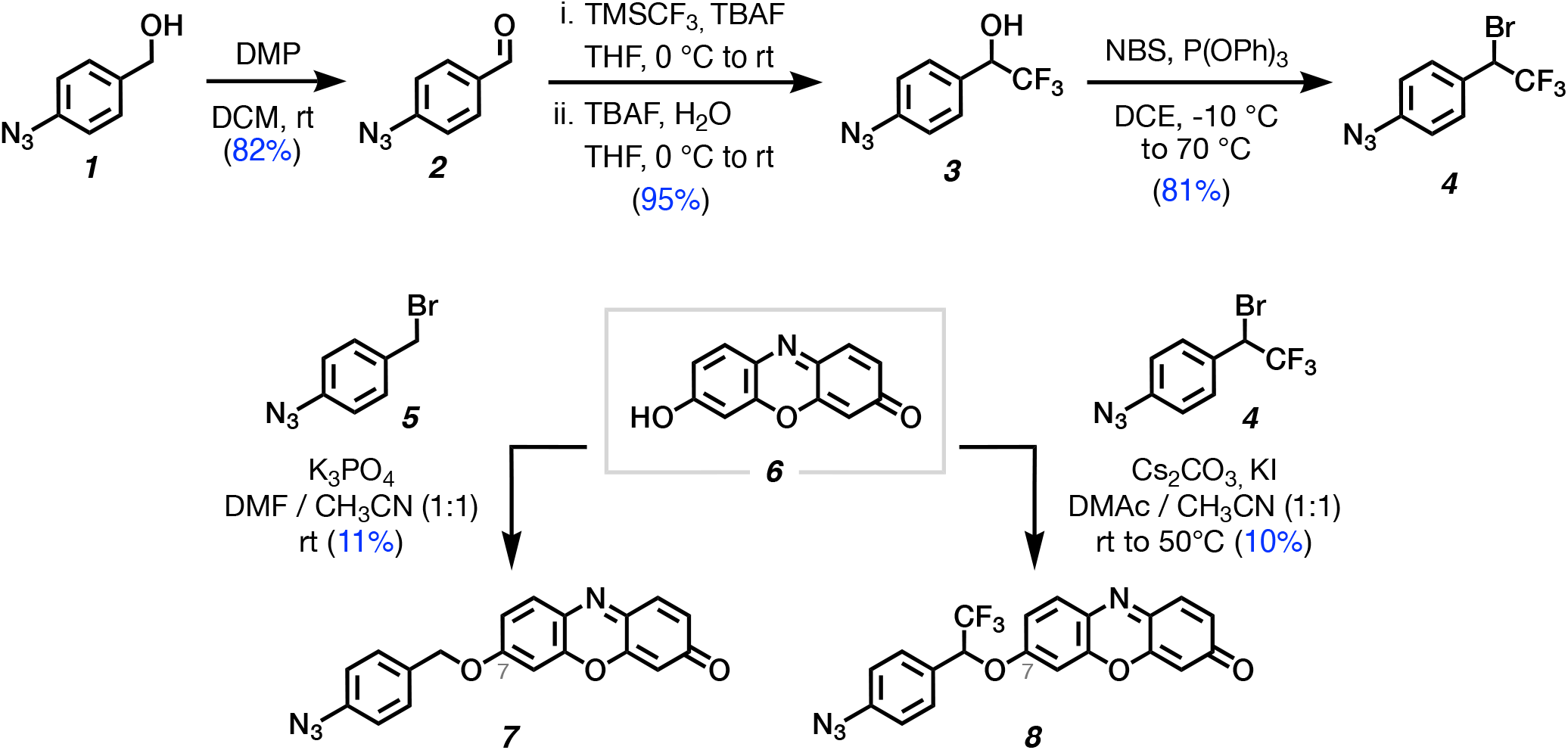
The chemical syntheses of the benzyl-caged resorufin *7* and α-CF_3_-benzyl-caged resorufin *8*.

### α-CF_3_ on 4-azidobenzyl group accelerates fluorescence generation and significantly improves H_2_S/HS^−^ detection sensitivity while maintaining high selectivity

Mammalian cell responses induced by hydrogen sulfide, such as angiogenesis, have been investigated using 10–600 μM exogenous H_2_S.^39^ In accordance, we reasoned that introducing 200 μM Na_2_S into a pH 7.5 medium (Tris, 100 mM) would produce H_2_S at a concentration that falls within this biologically relevant concentration range. At this pH, roughly 80% of the sulfur species is expected to exist in the form of HS^−^ (∼160 μM) and 20% in the form of H_2_S (∼40 μM), as the p*K*_a_ of H_2_S is ∼7.0.^40^ We consider that once administered into the aqueous buffer, virtually all of Na_2_S will be converted into its protonated forms, as the p*K*_a_ of HS^−^ is ≥12.^40^

We measured the fluorescence levels at 585 nm of the mixtures [***7*** and Na_2_S] and [***8*** and Na_2_S] following excitation at 545 nm. For each sample, we recorded the fluorescence intensities within one hour at 2-minute time intervals (Figures 3A and 3B). Upon addition of Na_2_S (200 μM estimated initial H_2_S/HS^−^ concentration), the signals from samples containing probe ***7*** barely (∼10%) increased within 1 hour incubation (Figure 3A), whereas those with ***8*** exhibited a steady increase, reaching to 2.6-fold intensity compared to the sample untreated (Figure 3B). These results indicate a significant acceleration in probe activation with the benzylic CF_3_ group in response to H_2_S/HS^−^, supporting our hypothesis that an electron withdrawing group incorporation at the benzylic position can assist probe release.

**Figure 3.**
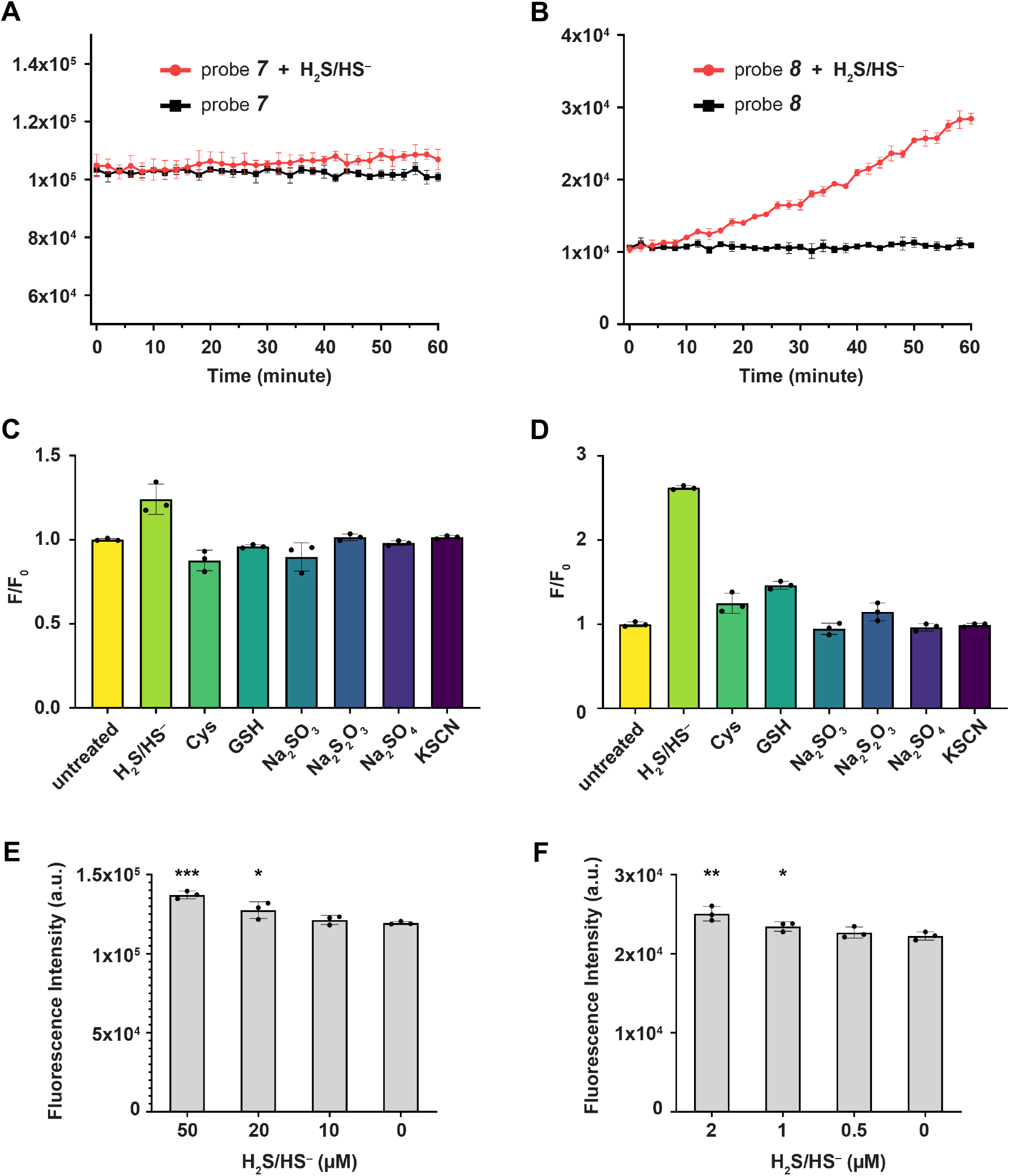
Probe performance in vitro. Activation kinetics **(A-B)**, Specificity **(C-D)** and limit of detection (LoD) **(E-F)** of ***7*** versus ***8***. LoD measurements were performed in Tris buffer (100 mM, pH 7.5). Error bars represent the standard deviation; n = 3. Single-tailed Student’s *t*-test: *, P < 0.05; **, P < 0.01; ***, P < 0.001.

Evaluation of the sulfur selectivity of the probes was conducted by using biologically relevant sulfur compounds: H_2_S/HS^−^, cysteine (Cys), glutathione (GSH), sulfite (SO_3_^2−^), sulfate (SO_4_^2−^), thiosulfate (S_2_O_3_^2−^), and potassium thiocyanate (KSCN). For an estimated sulfur species concentration of 200 µM for H_2_S/HS^-^ and 500 µM for others at the time of addition and for a total incubation time of 1 hour, *F/F*_*0*_ values were calculated for each species. Here, *F* represents the fluorescence intensity of the sample incubated with a defined sulfur species, while *F*_*0*_ is that of the untreated sample. The signals were measured following 545 nm excitation (see Supporting Information for details), and *F/F*_*0*_ for the untreated sample (control) was set to 1. For samples containing ***7*** (Figure 3C), the sulfur species other than H_2_S/HS^−^ induced an increase in fluorescence by less than 10%. This marginal signal enhancement is in agreement with the well-established hydrogen sulfide selectivity of the 4-azidobenzyl group.^41^ For those containing probe ***8*** (Figure 3D), the signal enhancement was less than 50%. The probe ***7*** provided ∼20% fluorescence enhancement upon treatment with H_2_S/HS^−^, while probe ***8*** delivered ≥2.6-fold enhancement, which constitutes a ∼16-fold increase compared to non-target sulfur species. These results indicate that installation of α-CF_3_ in the 4-azidobenzyl group improves the probe performance while the azide reduction remains highly selective for H_2_S/HS^−^ against the biologically relevant sulfur species we have tested.

Limit of detection (LoD) measurements enabled the assessment of how α-CF_3_ installation on the 4-azidobenzyl group influence H_2_S/HS^−^ detection sensitivity. We determined LoD for both ***7*** and ***8*** by evaluating the statistical differences between *F* and *F*_*0*_ upon 1 hour of incubation with varying concentrations of Na_2_S, where *F* and *F*_*0*_ are the fluorescence intensities of samples that were treated with Na_2_S and untreated, respectively. Here, LoD is defined as the minimum sulfur species concentration at which *F* is statistically higher than *F*_*0*_ using the one-tailed Student’s *t* test. Based on these parameters, the LoD for probe ***7*** was determined as 20 µM (Figure 3E), whereas that for probe ***8*** was 1 µM (Figure 3F). This significantly lower detection limit can be attributed to the increased sensitivity of ***8*** on its accelerated 1,6-elimination, which provides a larger number of fluorescently activated resorufin molecules at a given incubation time.

### DFT calculations indicate that benzylic CF_3_ substitution leads to a significantly lower activation barrier for the transition state of 1,6-elimination

To understand the impact of benzylic CF_3_ substitution on the energetics of 1,6-elimination, we performed DFT calculations using the Gaussian 16 program (Figure 4).^42^ The electronic structures were described with the B3LYP functional^43–45^ with D3 dispersion correction^46^ and 6-311++G(d,p) basis set. Additionally, the polarizable continuum model was applied with a dielectric constant of 78.4 to mimic the aqueous environment.^47^ For each reaction pathway, we conducted geometry optimization of both the reactants and products and determined their Gibbs free energies to obtain the reaction free energy, Δ*G*_*rxn*_ = *G*_*products*_ -*G*_*reactants*_. The synchronous transit-guided quasi-Newton method was used to search for the transition states,^48^ and the activation free energy was computed as Δ*G*_a_ = *G*_*TS*_ -*G*_*reactants*_. Vibrational frequency analysis was conducted to confirm that optimized geometries corresponded to the ground state stationary points or transition states. Equilibrium geometries exhibited solely real vibrational frequencies, while transition state structures showed a single imaginary frequency. Additional details on reaction paths and computational analyses are provided in the Supporting Information.

**Figure 4.**
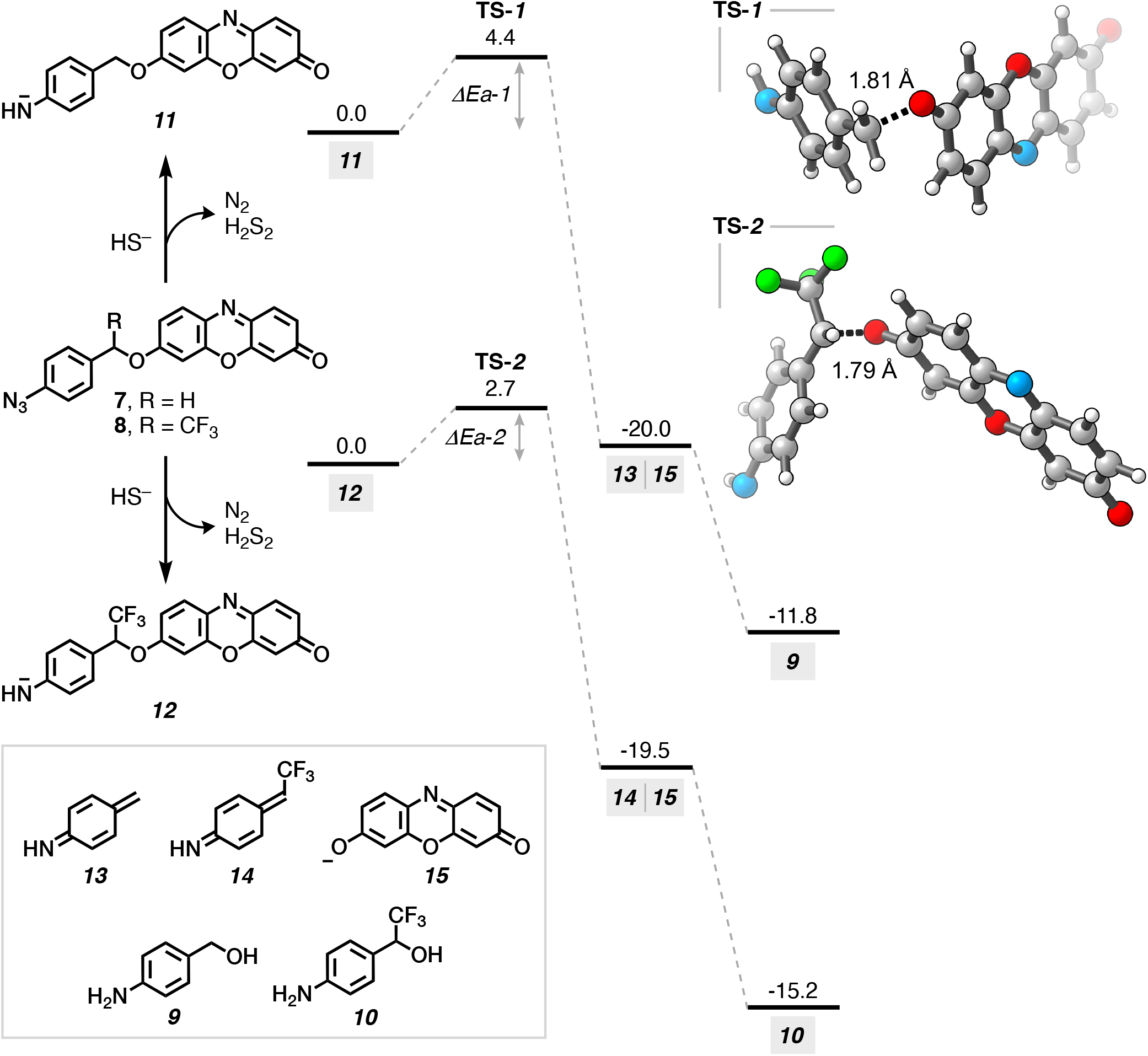
Computed free energy profiles for the 1,6-elimination of the caged resorufins *11* and *12*. All free energies were calculated under 1 atmospheric pressure and 298 K, and are reported in units of kcal/mol. Atomic representations in the molecular structures are C: grey, O: red, N: blue, F: green, and H: white.

Based on a combined experimental and computational investigation, Henthorn and Pluth reported that the reduction of 4-methyl-7-azidocoumarin to 4-methyl-7-aminocoumarin is mediated by two equivalents of HS^−^.^18^ In this reduction process, HS^−^ attacks on the terminal azide nitrogen of 4-methyl-7-azidocoumarin and forms a HS-(N=N)-NH-coumarin intermediate. Nucleophilic attack on this intermediate by the second HS^−^ generates N_2_, H_2_S_2_, and ^−^NH-coumarin anion, which then abstracts a proton intermolecularly, forming the neutral aminocoumarin (NH_2_-coumarin). Following their insightful work, we investigated how 1,6-elimination processes differ energetically for the azidobenzyl caged resorufins ***7*** and ***8***. Our DFT calculations showed that molecules ***11*** and ***12***, the putative aryl amide products to be generated from ***7*** and ***8***, can undergo an energetically favorable 1,6-elimination, each with a Δ*G*_rxn_ of approximately -20 kcal/mol (Figure 4). In contrast, the protonated forms of ***11*** and ***12*** (molecules ***19*** and ***20***, Figure S1) exhibit Δ*G*_rxn_ values of 10.8 kcal/mol and 16.0 kcal/mol, respectively (Table S1). The large and positive Δ*G*_rxn_ suggests that these 1,6-elimination reactions are unlikely to occur spontaneously at ambient conditions. The calculations also demonstrated a substantial difference in the activation free energies for the 1,6-elimination of ***11*** and ***12***. Their respective transition states, **TS-*1*** and **TS-*2***, involve the stretching of the ether C-O bond to ∼1.8 Å (Figure 4).

The strongly electron-withdrawing benzylic CF_3_ group in ***12*** stabilizes the negatively charged **TS-*2***, which results in a Δ*G*_*a*_ of 2.7 kcal/mol. This value is 1.7 kcal/mol lower than that for **TS-*1***, leading to a significant (17-fold) enhancement in the 1,6-elimination rate based on the Arrhenius equation. This difference in Δ*G*_*a*_ for **TS-*1*** vs **TS-*2*** aligns closely with the experimental observation that ***8*** displays a marked acceleration in fluorescence generation (*cf*. Figure 3B). 1,6-elimination of ***11*** and ***12*** produces the *para*-azaquinone-methides ***13*** and ***14***, respectively, along with ***15***, which is the aryloxide anion form resorufin (***6***). Hydration of these *para*-azaquinone-methides are highly favorable, as the formation of benzyl alcohol byproducts ***9*** and ***10*** is strongly exothermic (Δ*G*_rxn_ of -11.8 and -15.2 kcal/mol, respectively). From the DFT calculations, the benzylic CF3 substitution significantly enhances the rate of 1,6-elimination through stabilization of both the reaction transition states and products. These findings are consistent with the observed experimental reaction acceleration and provide important insights into the design of high-performing probes.

### The prevalence of intracellular hydrogen sulfide was captured in near-real-time

Certain cancer cells have been used to investigate intracellular H_2_S.^20,49,50^ We used HeLa as a mammalian cell model to investigate H_2_S/HS^−^ through activity-based sensing under controlled and stimulated conditions. Cell viability assay enabled us to assess the cytocompatibility of both 7-*O*-caged resorufins and determine a feasible probe concentration for live cell imaging. We employed 3-(4,5-dimethylthiazol-2-yl)-2,5-diphenyl tetrazolium bromide (MTT) in HeLa incubated with ***7*** or ***8*** at a concentration range of 4–250 µM over a 24-hour period (see Supporting Information for details). Dose-response curves (Figure S2) showed high IC_50_ values for each probe: ***7*** at 37 µM and ***8*** at 76 µM (Table S2). These results suggested that both probes do not induce cytotoxicity under our cell imaging conditions (20 µM, 30 minutes incubation).

Confocal fluorescence imaging of H_2_S/HS^−^ in HeLa was conducted as two different investigations. In the first study (Figure 5), we comparatively assessed the intracellular signal generation due to hydrogen sulfide that is either provided exogenously from media with Na_2_S addition or produced intracellularly following stimulation with L-Cys. In the second study (Figure 6), confocal imaging was performed to capture time-dependent prevalence of H_2_S/HS^−^ in HeLa upon stimulation with L-Cys.

**Figure 5.**
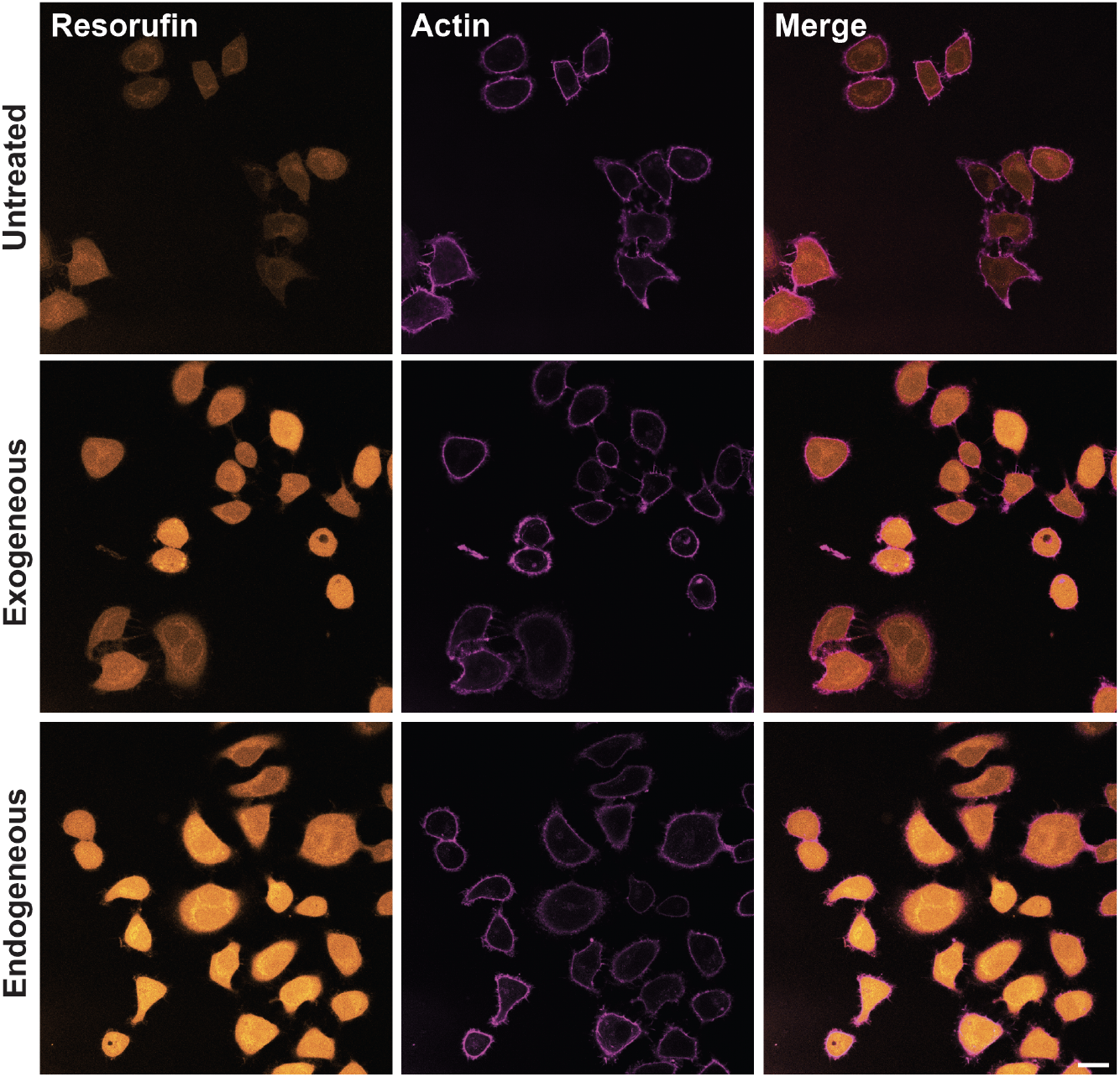
Confocal imaging of H_2_S/HS^−^ using probe *8* in live HeLa cells. Cells were incubated with probe ***8*** (20 μM) for 30 minutes and stained with actin dye (CellMask™ Deep Red Actin Tracking Stain) prior to imaging. **(Top row)** Cells were left untreated. **(Middle row)** Cells were treated with Na_2_S (200 μM) as the exogeneous source H_2_S/HS^−^. **(Bottom row)** Cells were treated with L-Cys (1 mM) as the substrate to induce endogenous generation of H_2_S/HS^−^. Cells were imaged after 1 hour following treatment with sulfur species. Resorufin channel: 571/583 nm; Actin channel: 652/669 nm. Scale bar: 20 µm.

**Figure 6.**
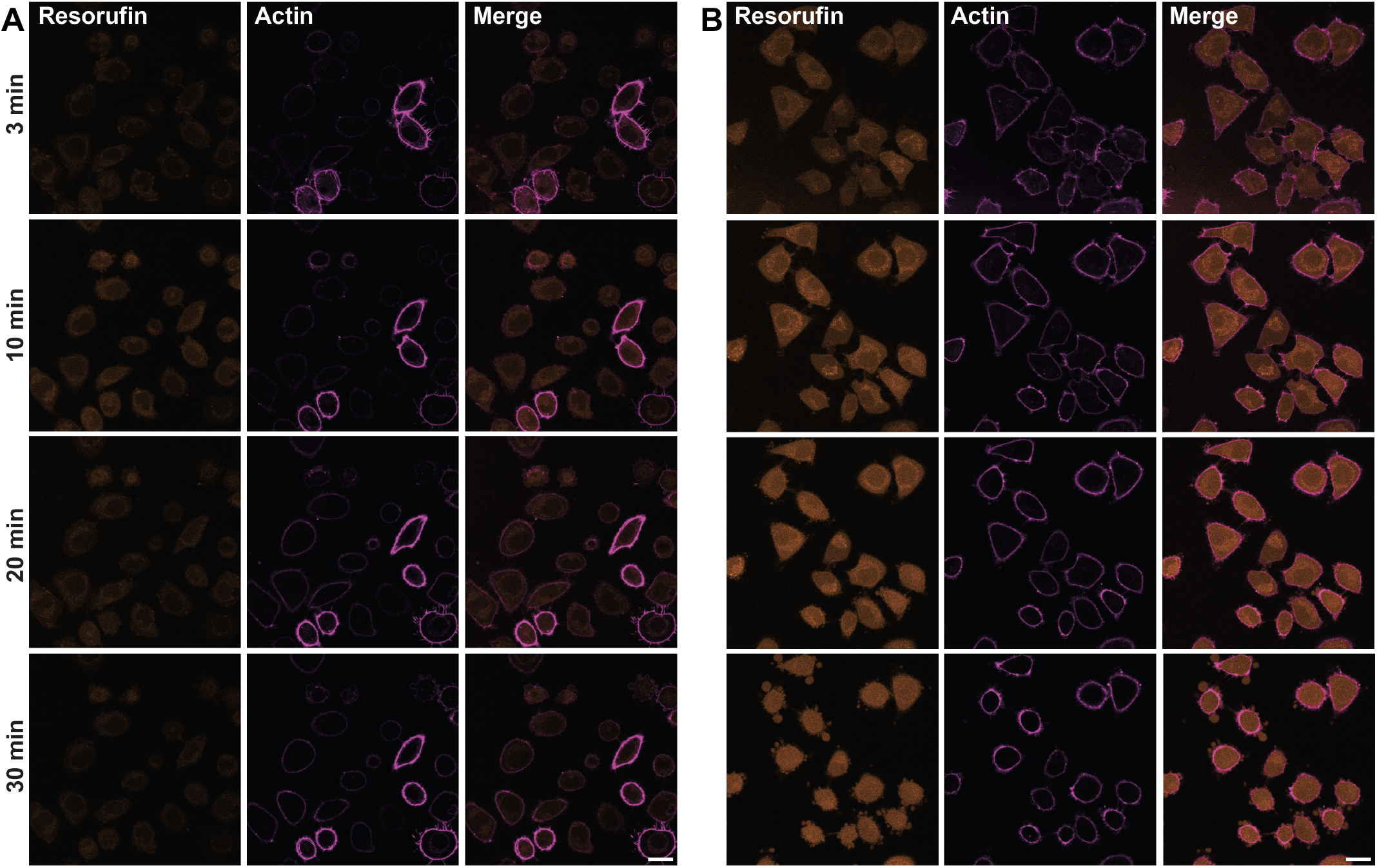
Confocal imaging of time-dependent capturing of H_2_S/HS^−^ generation in stimulated HeLa cells. Cells were incubated with either **(A)** probe ***7*** (20 μM) or **(B) *8*** (20 μM) and then stained with actin dye (CellMask™ Deep Red Actin Tracking Stain) prior to imaging. Cells were treated with L-Cys (1 mM) as a substrate to CSE and CBS to induce endogenous H_2_S/HS^−^. Apoptosis is visible in both images at the 10 min mark and necrosis of the majority of cells by 30 min. Resorufin channel: 571/583 nm; Actin channel: 652/669 nm. Scale bar: 20 µm.

HeLa were revived and seeded in DMEM (Dulbecco’s Modified Eagle Medium), 10% FBS (fetal bovine serum) and 100 U/mL penicillin. Actin staining served as a visual aid to highlight the cytoskeleton of the whole cell. With the probe ***8*** (20 μM in media), cells exhibited weak fluorescence without further treatment (Figure 5, top row), which can be associated with basal levels of intracellular hydrogen sulfide or the inherent fluorescence of ***8***, or more likely the combination of both. Notably, those incubated with ***8*** followed by 200 µM Na_2_S showed significant fluorescence signal enhancement, indicative of an efficient probe turn-on process due to a greater level of H_2_S/HS^−^. (Figure 5, middle row). Cells were also incubated in L-Cys (1 mM) to stimulate endogenous synthesis of H_2_S/HS^−^, which produced the highest increase of fluorescence (Figure 5, bottom row) in comparison to the untreated samples. The increase of signal in the presence of either Na_2_S or L-Cys through the use of ***8*** indicates proper detection of hydrogen sulfide intracellularly within live HeLa cells.

To validate the intracellular kinetics of ***7*** and ***8***, a time-based study was conducted with HeLa using L-Cys (1 mM in HBSS) (Figure 6). HeLa cells were incubated with either probes ***7*** or ***8*** at 20 μM, and confocal images were acquired at five time points (3, 10, 20, 30, and 40 minutes) post stimulation. Images indicated the start of apoptosis within 10 minutes following stimulation with L-Cys, and majority of cells were necrotic after 30 minutes. This observation is consistent with the previously demonstrated changes in morphology of HeLa under stress due to a topoisomerase inhibitor.^51^ Notably, we observed a significant qualitative difference in signal between the cells with different probes (Figure 6A versus 6B): those containing ***8*** exhibited greater levels of fluorescence enhancement. The intracellular fluorescence (resorufin channel) was observable in just 3 minutes post L-Cys addition, the earliest possible image acquisition time upon cell treatment in our imaging setup. This indicated that, within the untreated cells existing H_2_S/HS^−^ can lead to the activation of probe **8**, which is consistent with the observation in Figure 5 (top row). The change in signal intensity was distinguishable until ∼20 minutes, after which the cells were observed to progressively lose their cytoskeletal integrity (30-minute time point, Figure 6; also see Figure S3).

### CONCLUSION

In conclusion, we have discovered a highly efficient method for fluorescent probe activation, which is driven by H_2_S-triggered 1,6-elimination of α-CF_3_-benzyl caging group. Activity-based sensing utilizes the innate reactivity of caged fluorogenic molecules to detect analytes, and 1,6-elimination of benzyl groups is a mechanism commonly implemented in probe activation strategies. Rapid response rate and high target sensitivity are probe activation criteria that are challenging to achieve, especially in the case of detecting RSS by confocal microscopy. Our α-CF_3_-benzylation approach can markedly enhance these properties while maintaining selectivity. 4-azido-(α-CF_3_)-benzyl resorufin (probe ***8***) compared to its protio analog (***7***) offers significantly faster signal generation and improved sensitivity in response to H_2_S.

DFT calculations support the hypothesis that a benzylic CF_3_ group can reduce the activation energy barrier for 1,6-elimination. Live cell imaging demonstrates the superior performance of ***8***, with which we can now detect the prevalence of intracellular H_2_S in near-real-time. Our concurrent efforts will demonstrate that the described approach is translatable to applications beyond bioimaging, such as the medicinal chemistry of prodrugs^52^ and antibody-drug conjugates.^53^

### SUPPORTING INFORMATION

The Supporting Information includes chemicals and materials; synthesis protocols and characterizations of compounds; method details for cell work; and supplementary tables and figures.

## ACKNOWLEDGMENT

We thank Mark J. Dresel and Dorsa Ebrahimi for useful discussions. We thank Dr. Ki-Bum Lee for the donation of HeLa cells and Dr. Alan Goldman for the donation of triphenyl phosphite. We acknowledge the Office of Advanced Research Computing at Rutgers University for providing access to the Amarel server.

## AUTHOR CONTRIBUTIONS

Liming W., A.S., and E.C.I. conceived the project. Liming W., A.S., and Y.K. conducted synthetic chemistry experiments and characterized novel compounds. S.C., Liming W., and T.A. conducted cell work. S.C. and Liming W. performed confocal imaging. R.Z. and Lu W. ran and analyzed DFT calculations. All of the authors contributed to the interpretation of data. E.C.I. oversaw the study and wrote the manuscript with input from all authors. CRediT: **Liming Wang** conceptualization, data curation, formal analysis; **Aditya Sivakumar** conceptualization, data curation, formal analysis; **Sarah Cho** data curation, formal analysis; **Rui Zhang** data curation, formal analysis; **Yuhyun Kim** data curation; **Tushar Aggarwal** data curation; **Lu Wang** formal analysis, resources, supervision; **Enver Cagri Izgu** conceptualization, formal analysis, project administration, resources, supervision, writing original draft.

