## Supporting Information for "Benzylic Trifluoromethyl Accelerates 1,6-Elimination Toward Rapid Probe Activation"

[§]These authors contributed equally

**Contents** **Page**


| Contents | Page |
|---|---|
| **Chemicals** | **S3** |
| **General Synthetic Methods and Instrumental Analyses** | **S4** |
| **Preparation and Characterization of Small Molecules** | **S5–S21** |
| Compound **2** | S5 |
| Compound **3** | S6 |
| Compound **4** | S7 |
| Compound **7** | S8 |
| Compound **8** | S9 |
| $^1$H, $^{13}$C, $^{19}$F NMR Spectra | S10–S21 |
| **Computational Methods and Results** | **S22** |
| Figure S1 | S23 |
| Table S1 | S23 |
| **Fluorescence Measurement** | **S24** |
| **Cell Studies** | **S25** |
| Figure S2 | S26 |
| Table S2 | S26 |
| **Impact of Endogenous Stimulation with L-Cys on HeLa** | **S27** |
| **References** | **S28** |

### Chemicals

**Reagents:** Sodium bicarbonate ($NaHCO_3$), sodium sulfate ($Na_2SO_4$), *N*-bromosuccinimide (NBS), triphenylphosphite ($P(OPh)_3$), cesium carbonate ($Cs_2CO_3$), potassium phosphate ($K_3PO_4$), hydrochloric acid (HCl), sodium nitrate ($NaNO_2$), sodium azide ($NaN_3$), potassium iodide (KI), sodium sulfide nonahydrate ($Na_2S{\cdot}9H_2O$) were purchased from Millipore-Sigma. 4-aminobenzyl alcohol, Dess-Martin periodinane (DMP), (trifluoromethyl)trimethylsilane ($TMSCF_3$) were purchased from Combi-Blocks. Tetrabutylammonium fluoride (TBAF) solution (1 M in THF) was purchased from Tokyo Chemical Industry.

**Solvents, Buffers, Media:** Acetone, dichloromethane ($CH_2Cl_2$), 1,2-dichloroethane (DCE), dimethylformamide (DMF), ethyl acetate (EtOAc), *N*,*N*−dimethylacetamide (DMAc), and tetrahydrofuran (THF) were purchased from Millipore Sigma. Acetonitrile ($CH_3CN$), diethyl ether, hexanes, and methanol (MeOH, HPLC grade) were purchased from Fisher Scientific. Deuterated solvents, containing 0.05% (v/v) TMS, were purchased from either Cambridge Isotope Laboratories. Water was deionized and filtered to a resistivity of 18.2 MΩ·cm with a Milli-Q® Plus water purification system (Millipore, Massachusetts). Phosphate-buffered saline (PBS), tris(hydroxymethyl)-aminomethane (Tris) were purchased from Millipore Sigma. Buffers were prepared freshly in Milli-Q® water and their pH were adjusted using HCl or NaOH using a Thermo Scientific Orion Star pH meter. Hanks' Balanced Salt Solution (HBSS) was purchased from VWR International.

### General Synthetic Methods and Instrumental Analyses

All reactions were performed under a dry nitrogen atmosphere unless otherwise stated. All glassware (e.g., round-bottom flasks, vials) was oven-dried before use. Purification of the synthesized compounds was performed using a Büchi Reveleris® flash chromatography system equipped with a silica (50 μm irregular) column. Nuclear magnetic resonance (NMR) spectroscopic analyses were carried out using a Bruker Avance Neo 500 MHz spectrometer. NMR data is provided for new compounds. $^{1}$H NMR spectra were acquired at 500 MHz, $^{13}$C NMR spectra were acquired at 126 MHz, and $^{19}$F NMR spectra were acquired at 471 MHz. Chemical shifts ($\delta$) for $^{1}$H NMR spectra were referenced to $(C\boldsymbol{H}_3)_4Si$ at $\delta$ = 0.00 ppm or to $C\boldsymbol{H}Cl_3$ at $\delta$ = 7.26 ppm. $^{13}$C NMR spectra were referenced to $\boldsymbol{C}DCl_3$ at $\delta$ = 77.23 ppm or to $(\boldsymbol{C}H_3)_4Si$ at $\delta$ = 0.00 ppm.

The following abbreviations are used to describe $^{1}$H NMR resonances: s (singlet), d (doublet), t (triplet), q (quartet), m (multiplet), and dd (doublet of doublets). Coupling constants (*J*) are reported in Hz. Liquid chromatography followed by high-resolution mass spectrometry (LC-HRMS) analysis from electrospray ionization (ESI) was carried out on a Waters Acquity-Xevo G2-XS QTof instrument. Fluorescence measurements were conducted using a Spectra Max ID3 instrument (Molecular Devices) in Greiner 96-well half area black plates. Confocal fluorescence microscopy was performed on a Leica TCS SP8 tauSTED microscope with a 40” objective using Leica type F immersion liquid. All statistical analyses were performed using GraphPad Prism software version 9.

### Preparation and Characterization of Small Molecules

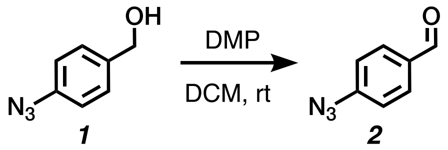


**Synthesis of 4-azidobenzaldehyde (2):** 4-azidobenzyl alcohol (**1**) was synthesized from 4-aminobenzyl alcohol following a previously reported one-step protocol.[1] To 100-mL round-bottom flask containing a vigorously stirring solution of **1** (535 mg, 3.59 mmol, 1.0 equiv) in $CH_2Cl_2$ (24 mL), Dess-Martin periodinane (DMP) (3.046 g, 7.18 mmol, 2.0 equiv) was added. The reaction mixture was allowed to stir at room temperature (rt) and monitored by TLC until completion (3 hour). The resulting mixture was concentrated under reduced pressure and then re-dissolved in EtOAc. The organic solution was washed three times with aqueous $Na_2SO_3$ (15% w/w) through vigorous stirring (for 10 minutes), then washed with saturated aqueous $NaHCO_3$ twice. The organic layers were collected and dried with $Na_2SO_4$. Volatiles were removed under reduced pressure, providing **2** (435 mg, 82%) as a light yellow solid with a typical benzaldehyde odor. The product was at sufficient purity by NMR spectroscopy assessment and, thus, was used without further purification.

**$^1$H NMR** (500 MHz, $CDCl_3$): $\delta$ 9.93 (s, 1H), 7.87 (d, *J* = 8.5 Hz, 2H), 7.15 (d, *J* = 8.6 Hz, 2H).

**$^{13}$C NMR** (126 MHz, $CDCl_3$): $\delta$ 190.60, 146.29, 133.25, 131.56, 119.49.

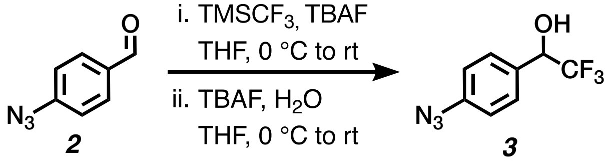


**Synthesis of 1-(4-azidophenyl)-2,2,2-trifluoroethan-1-ol (3):** To a 20-mL vial was added the aldehyde **2** (435 mg, 2.96 mmol, 1.00 equiv) in THF (10 mL) under a nitrogen-rich atmosphere. The mixture was cooled to 0 °C and $TMSCF_3$ (875 μL, 5.92 mmol, 2.00 equiv) was added, followed by the dropwise addition of TBAF (1 M in THF, 44 μL, 0.044 mmol, 0.015 equiv). The vial was then sealed, and the reaction mixture was allowed to stir at rt overnight. The mixture was then cooled to 0 °C, and $H_2O$ (293 μL, 16.27 mmol, 5.50 equiv) was added, followed by the dropwise addition of TBAF (1 M in THF, 296 μL, 0.296 mmol, 0.10 equiv). The reaction mixture was stirred at rt overnight and then concentrated under reduced pressure. The resulting material was redissolved in diethyl ether and washed with water (40 mL) three times. The organic layer was collected, dried with $Na_2SO_4$, and concentrated under vacuum. The crude product was dissolved in acetone, and the volatiles were removed again under reduced pressure. This process was repeated twice more, which ultimately provided the alcohol **3** (610 mg, 95%) as a pale yellow solid. The product was at sufficient purity by NMR spectroscopy assessment and, thus, was used without further purification.

**$^1$H NMR** (500 MHz, $CDCl_3$): δ 7.47 (d, *J* = 8.6 Hz, 2H), 7.07 (d, *J* = 8.6 Hz, 2H), 5.02 (q, *J* = 6.6 Hz, 1H), 2.71 (s, 1H).

**$^{13}$C NMR** (126 MHz, $CDCl_3$): δ 141.40, 130.43, 128.99, 124.12 (q, *J* = 282.1 Hz), 119.20, 72.28 (q, *J* = 32.2 Hz).

**$^{19}$F NMR** (470 MHz, $CDCl_3$): δ -78.54 (d, *J* = 6.7 Hz).

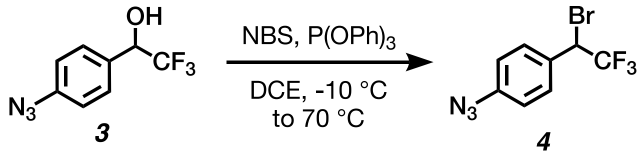

**Synthesis of 1-azido-4-(1-bromo-2,2,2-trifluoroethyl)benzene (compound 4):** To a 250-mL round-bottom flask was loaded a solution of NBS (1.2 g, 6.74 mmol, 2.4 equiv) in DCE (30 mL) under a nitrogen-rich atmosphere. The solution was cooled to -10 °C while being stirred vigorously, and triphenylphosphite (1.476 mL, 5.61 mmol, 2.0 equiv) was added dropwise. After 5 minutes of mixing, a solution of the alcohol **3** (610 mg, 2.81 mmol, 1.0 equiv) in DCE (10 mL) was added to flask dropwise. The resulting mixture was warmed to 70 °C, at which it was stirred until completion (~6 hours) based on $^{19}$F NMR spectroscopic analysis. The reaction mixture was then cooled to rt, concentrated under reduced pressure, and redissolved in diethyl ether. The solution was washed twice with 0.5 M aqueous $H_2SO_4$. The organic layer was collected, dried with $Na_2SO_4$, and concentrated under reduced pressure to an oil. This crude product was purified by silica gel column chromatography (mobile phase: EtOAc / hexanes, step gradient from 0 to 5% EtOAc). The product fractions were collected at 0% EtOAc and concentrated under reduced pressure, providing the benzyl bromide **4** as a dense yellow oil (635 mg, 81%).

**$^{1}$H NMR** (500 MHz, $CDCl_3$): δ 7.50 (d, *J* = 8.6 Hz, 2H), 7.05 (d, *J* = 8.7 Hz, 2H), 5.11 (q, *J* = 7.3 Hz, 1H).

**$^{13}$C NMR** (126 MHz, $CDCl_3$): δ 141.93, 130.68, 129.28, 123.30 (q, *J* = 278.0 Hz), 119.46, 46.44 (q, *J* = 34.4 Hz).

**$^{19}$F NMR** (470 MHz, $CDCl_3$): δ -70.56 (d, *J* = 7.3 Hz).

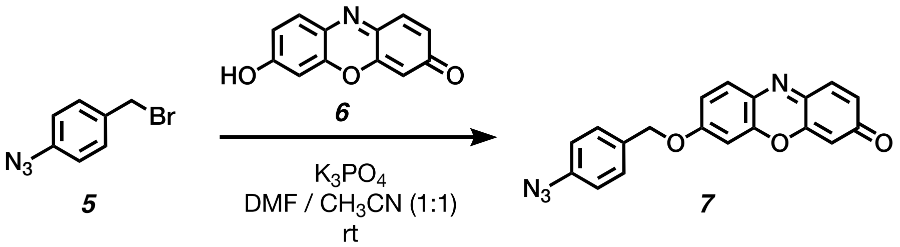


**Synthesis of 7-((4-azidobenzyl)oxy)-3*H*-phenoxazin-3-one (8):** 4-Azidobenzyl bromide **(7)** was synthesized from 4-aminobenzyl alcohol following a previously reported two-step protocol.[1] To a stirred solution of **7** (111.0 mg, 0.523 mmol, 1.0 equiv) in 10.5 mL of DMF/$CH_3CN$(1:1, v/v) under a nitrogen-rich atmosphere was added **6** (167.9 mg, 0.785 mmol, and 1.5 equiv), and $K_3PO_4$ (166.6 mg, 0.785 mmol, 1.5 equiv). After being stirred for 3 days at room temperature, the reaction mixture was concentrated under reduced pressure, and the crude product was purified by flash column chromatography with a neutral alumina column (mobile phase: $CH_2Cl2$ / MeOH, step gradient from 0 to 15% MeOH). The product fractions were collected and concentrated under reduced pressure, providing a crude mixture of **7**. The crude product was subsequently triturated with EtOAc thrice to provide **8** (20.4 mg, 11%) as an orange solid

**$^1$H NMR** (500 MHz, $CDCl_3$): δ 7.72 (d, *J* = 8.9 Hz, 1H), 7.46 – 7.39 (m, 3H), 7.11 – 7.06 (m, 2H), 7.00 (dd, *J* = 9.0, 2.7 Hz, 1H), 6.89 – 6.81 (m, 2H), 6.32 (t, *J* = 2.0 Hz, 1H), 5.14 (s, 2H).

**$^{13}$C NMR** (126 MHz, $CDCl_3$): δ 186.31, 162.41, 149.78, 145.85, 145.60, 140.45, 134.72, 134.33, 132.04, 131.66, 129.19, 128.56, 119.44, 114.18, 106.82, 101.08, 70.29.

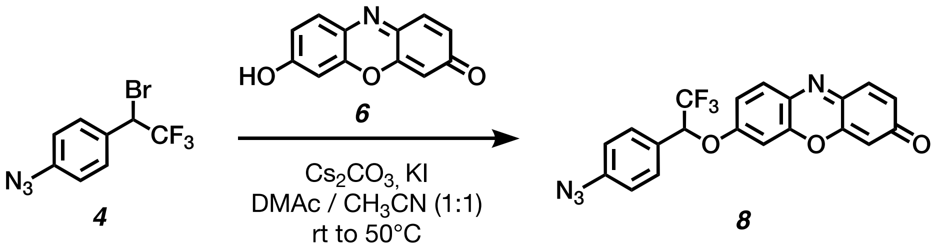


**Synthesis of 7-(1-(4-azidophenyl)-2,2,2-trifluoroethoxy)-3*H*-phenoxazin-3-one (5):** To a 1-dram vial was added a solution of **4** (20.0 mg, 0.071 mmol, 1.0 equiv) in 700 µL of DMAc/$CH_3CN$(1:1, v/v) under a nitrogen-rich atmosphere. To this solution was added **6** (22.8 mg, 0.107 mmol, and 1.5 equiv), $Cs_2CO_3$ (34.9 mg, 0.107 mmol, 1.5 equiv), and KI (11.9mg, 0.071 mmol, 1.0 equiv). After being stirred for 24 hours at room temperature, the reaction mixture was warmed to 50 °C and stirred for a further 24 hours. The reaction mixture was then cooled to rt and concentrated under reduced pressure. The crude product was purified by silica gel column chromatography (mobile phase: 30% EtOAc in hexanes), The product fractions were collected and concentrated under reduced pressure, delivering **8** (3 mg, ~10%) as an orange solid.

**$^1$H NMR** (500 MHz, $CDCl_3$): δ 7.69 (d, *J* = 8.9 Hz, 1H), 7.51 (d, *J* = 8.6 Hz, 2H), 7.39 (d, *J* = 9.8 Hz, 1H), 7.11 (d, *J* = 8.6 Hz, 2H), 6.96 (dd, *J* = 8.9, 2.7 Hz, 1H), 6.88 – 6.80 (m, 1H), 6.75 (d, *J* = 2.7 Hz, 1H), 6.28 (d, *J* = 2.0 Hz, 1H), 5.48 (q, *J* = 6.0 Hz, 1H).

**$^{19}$F NMR** (470 MHz, $CDCl_3$): δ -76.68 (d, *J* = 6.1 Hz)..

**HRMS** (ESI) m/z: Calculated for $C_{20}H_{12}F_3N_4O_3^+$ $[M + H]^+$ requires 413.0856; found 413.0854.

Compound ***2*** ($^1H$ NMR: 500 MHz, $CDCl_3$)

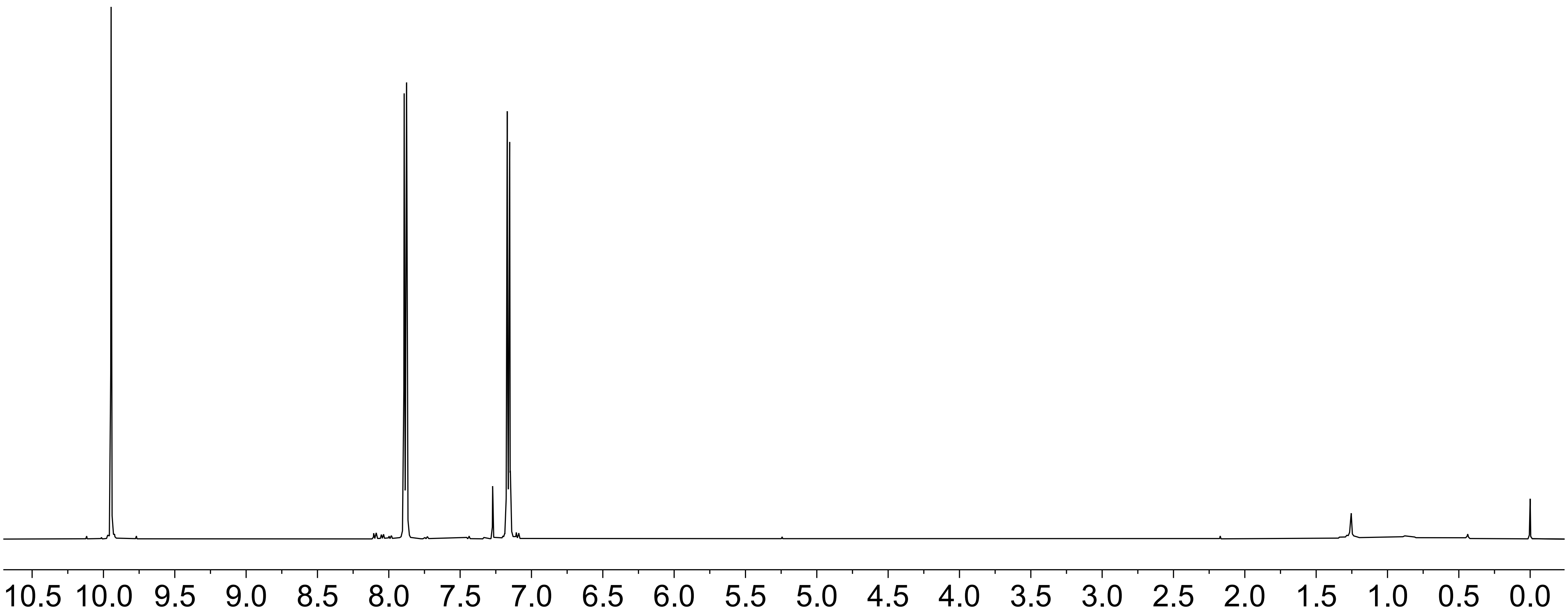

Compound ***2*** ($^{13}$C NMR: 126 MHz, $CDCl_3$)

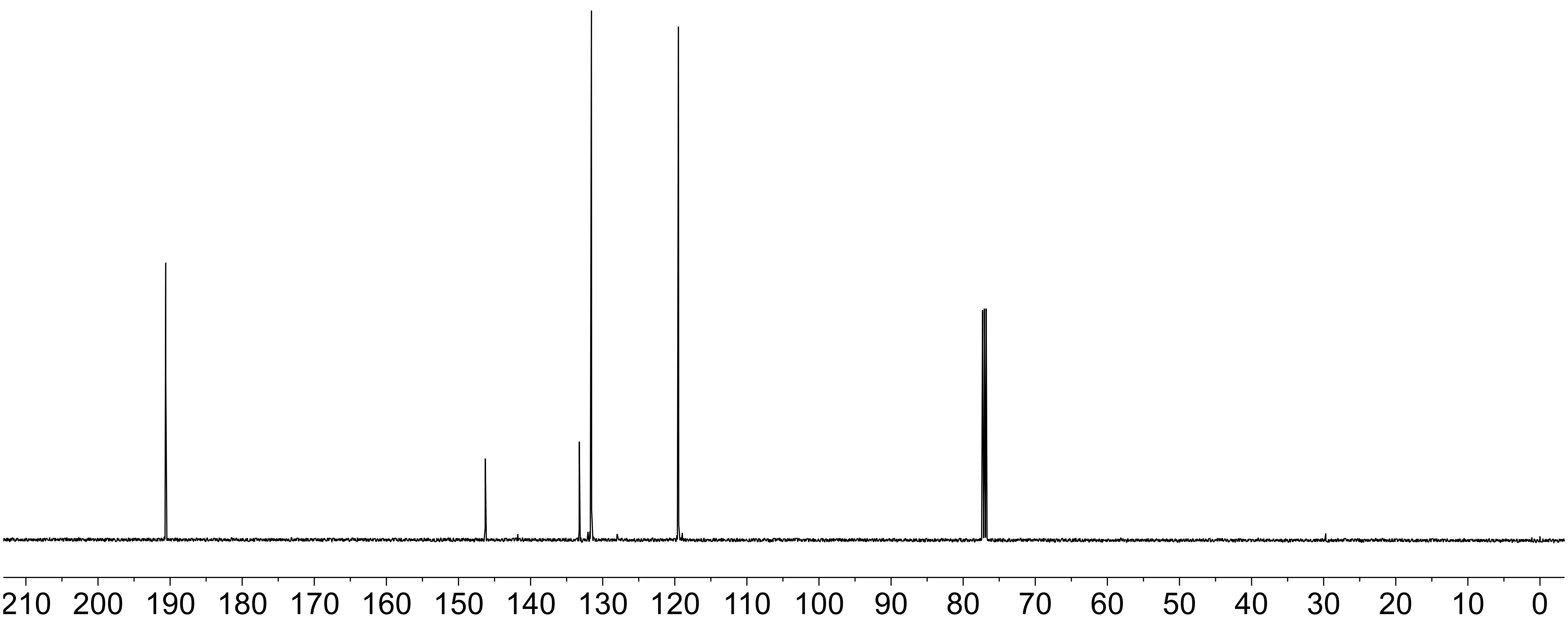

Compound ***3*** ($^1$H NMR: 500 MHz, $CDCl_3$)

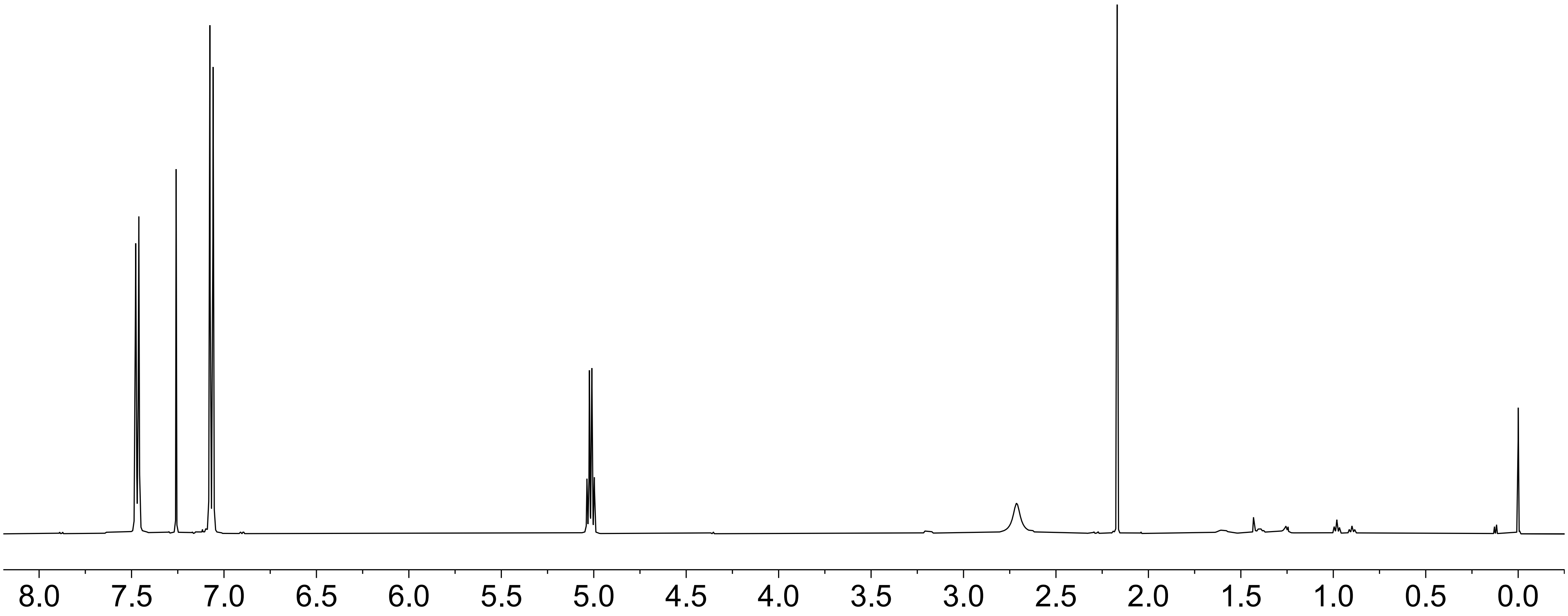

Compound ***3*** ($^{13}C$ NMR: 126 MHz, $CDCl_3$)

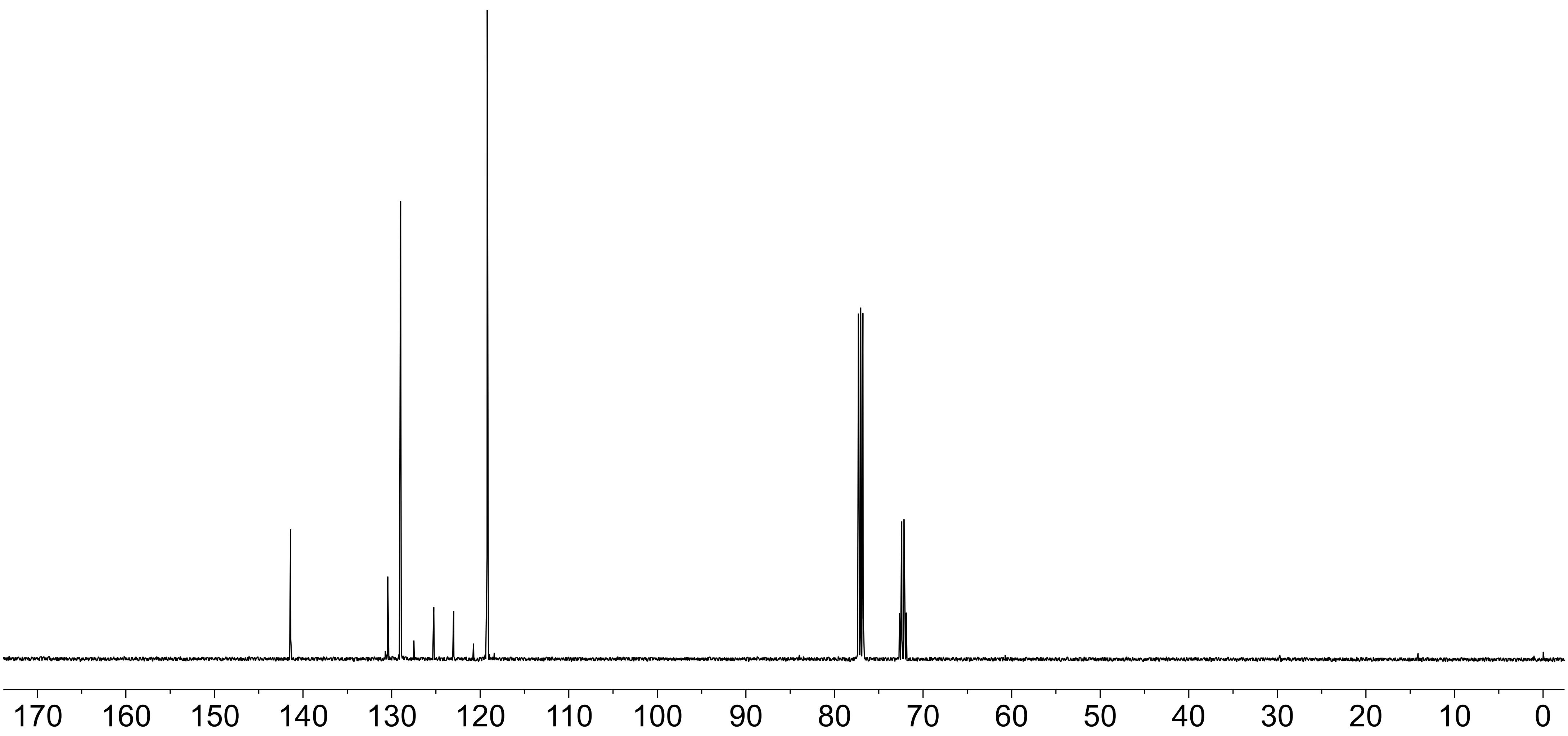

Compound ***3*** ($^{19}$F NMR: 470 MHz, $CDCl_3$)

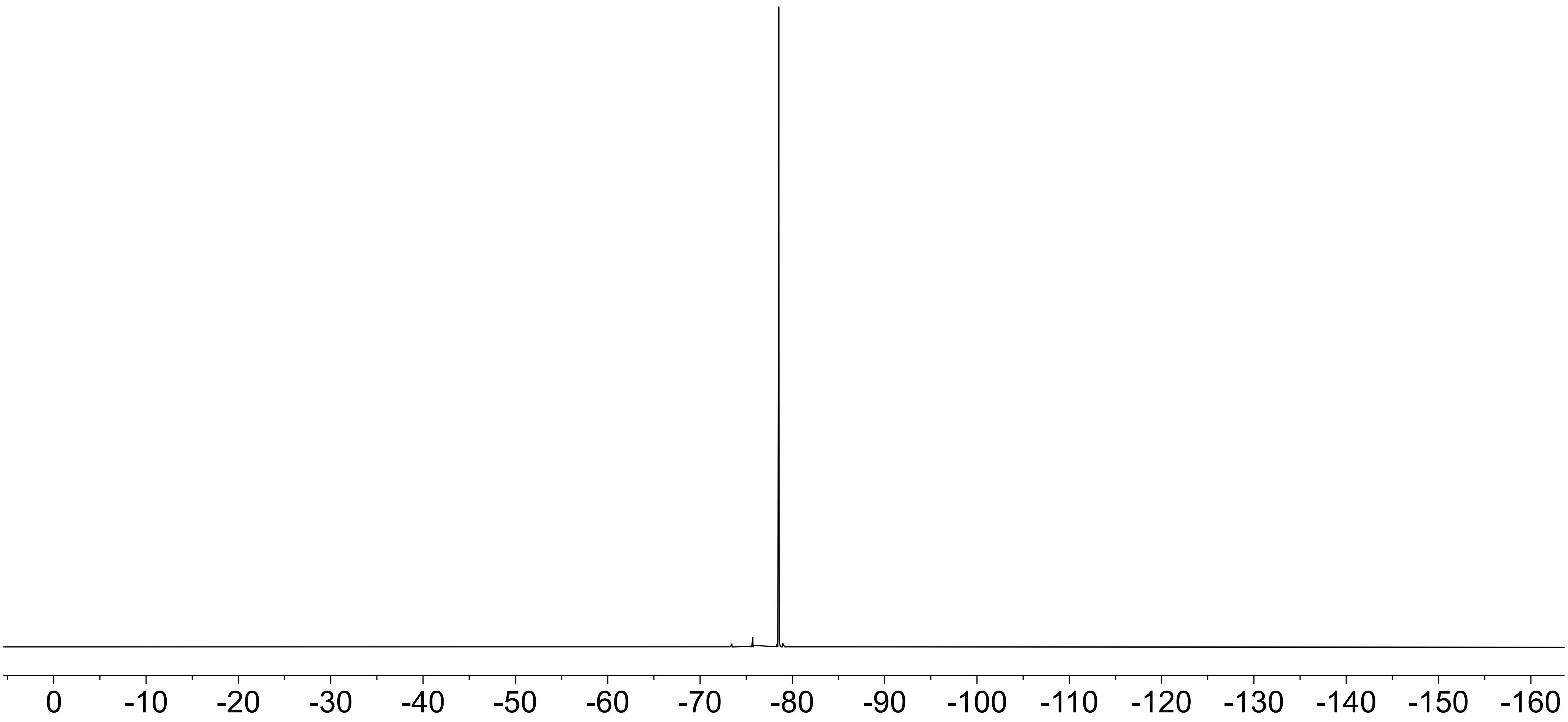

Compound ***4*** ($^1$H NMR: 500 MHz, $CDCl_3$)

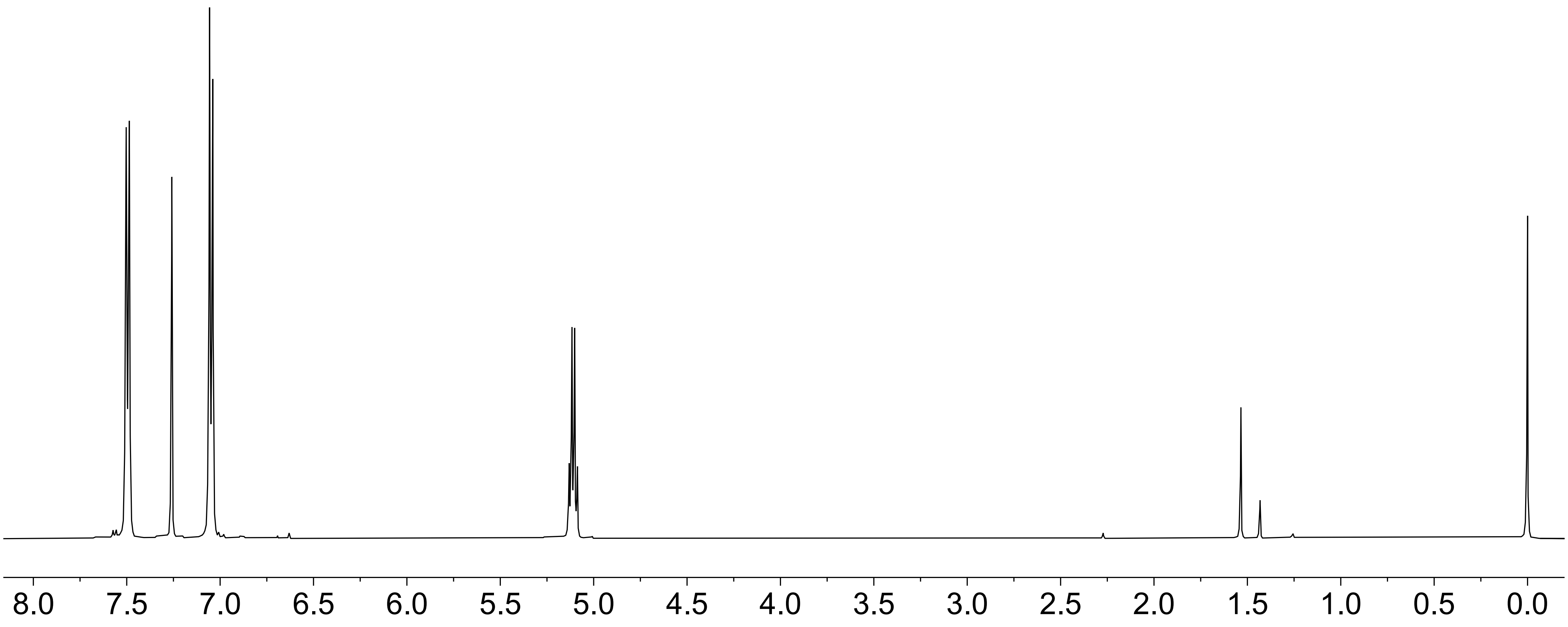

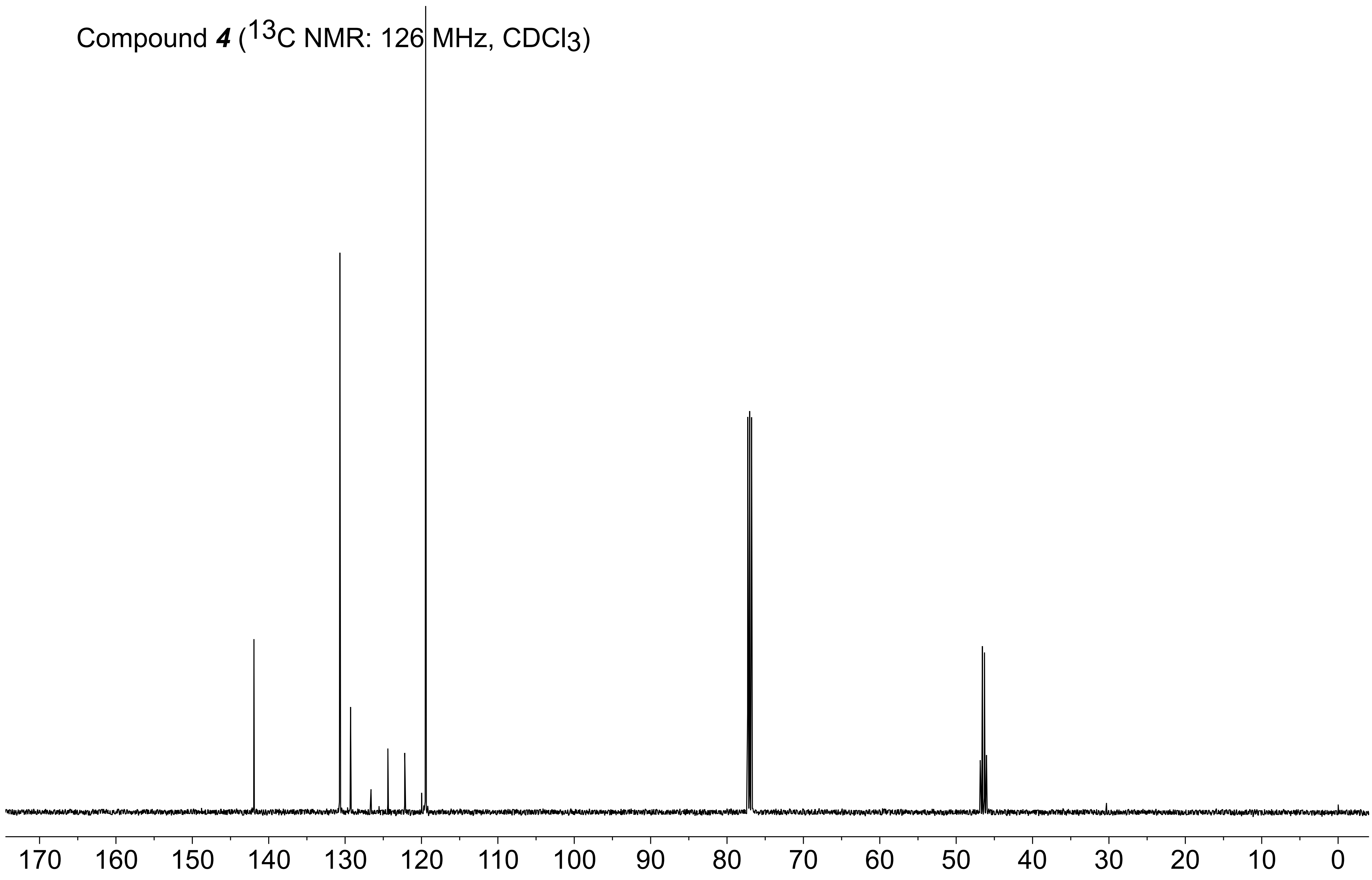
Compound 4 (13C NMR: 126 MHz, CDCl3)
170 160 150 140 130 120 110 100 90 80 70 60 50 40 30 20 10 0

Compound ***4*** ($^{19}$F NMR: 470 MHz, $CDCl_3$)

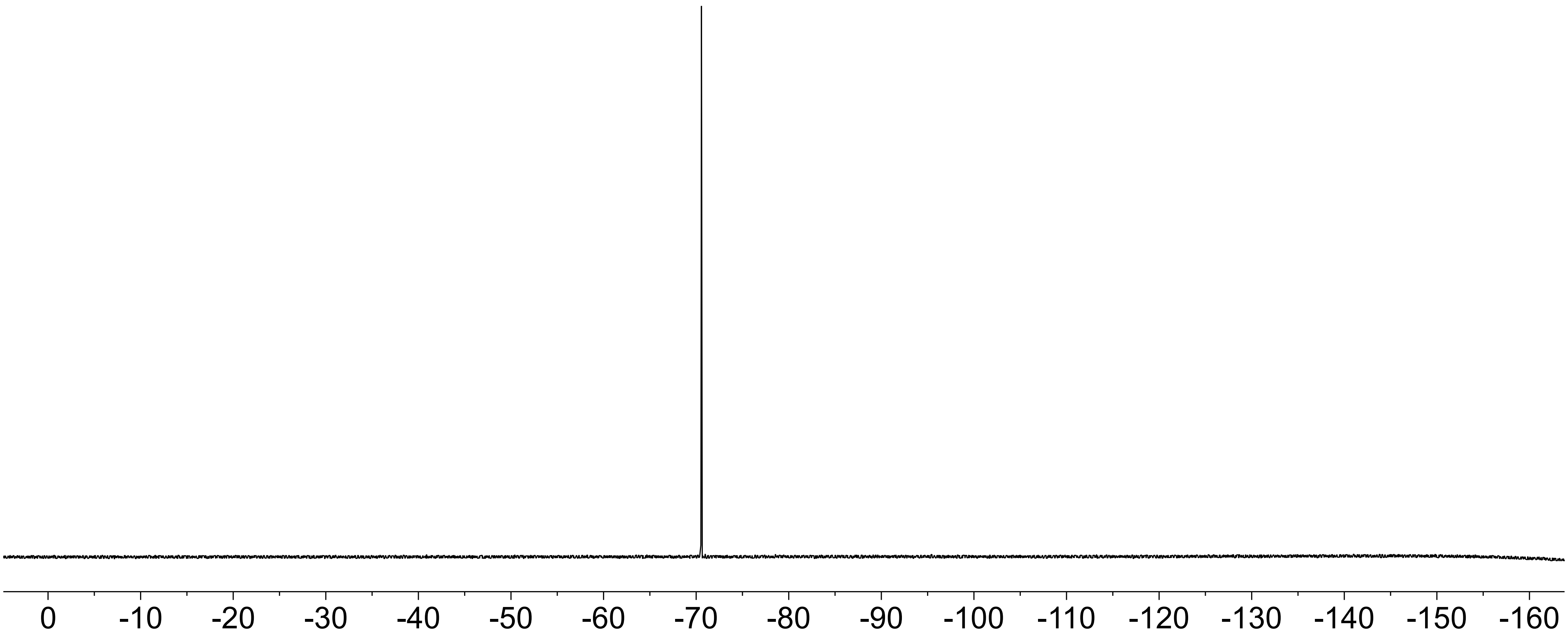

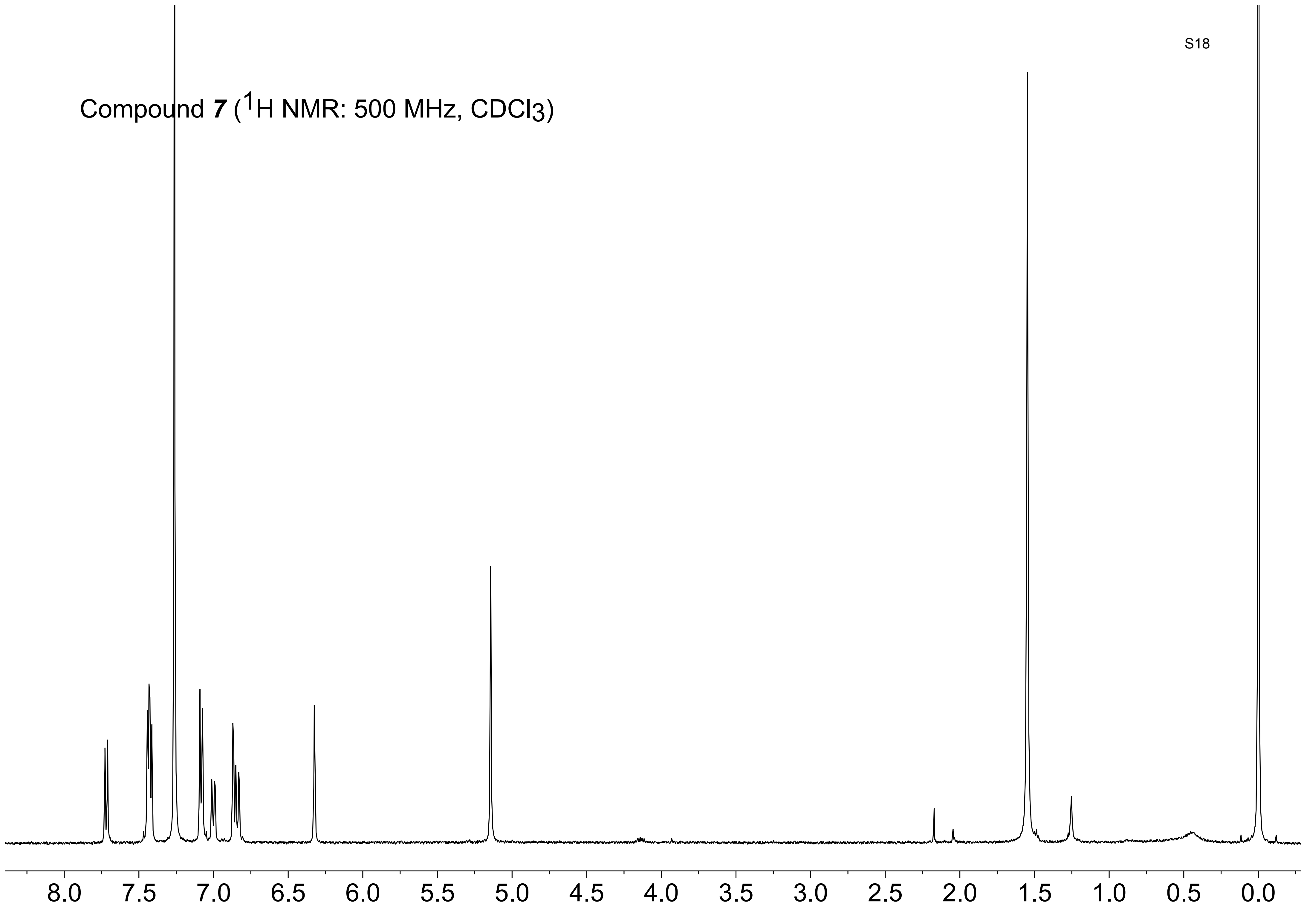
Compound **7** ($^1$H NMR: 500 MHz, $CDCl_3$)
8.0
7.5
7.0
6.5
6.0
5.5
5.0
4.5
4.0
3.5
3.0
2.5
2.0
1.5
1.0
0.5
0.0

Compound **7** ($^{13}C$ NMR: 126 MHz, $CDCl_3$)

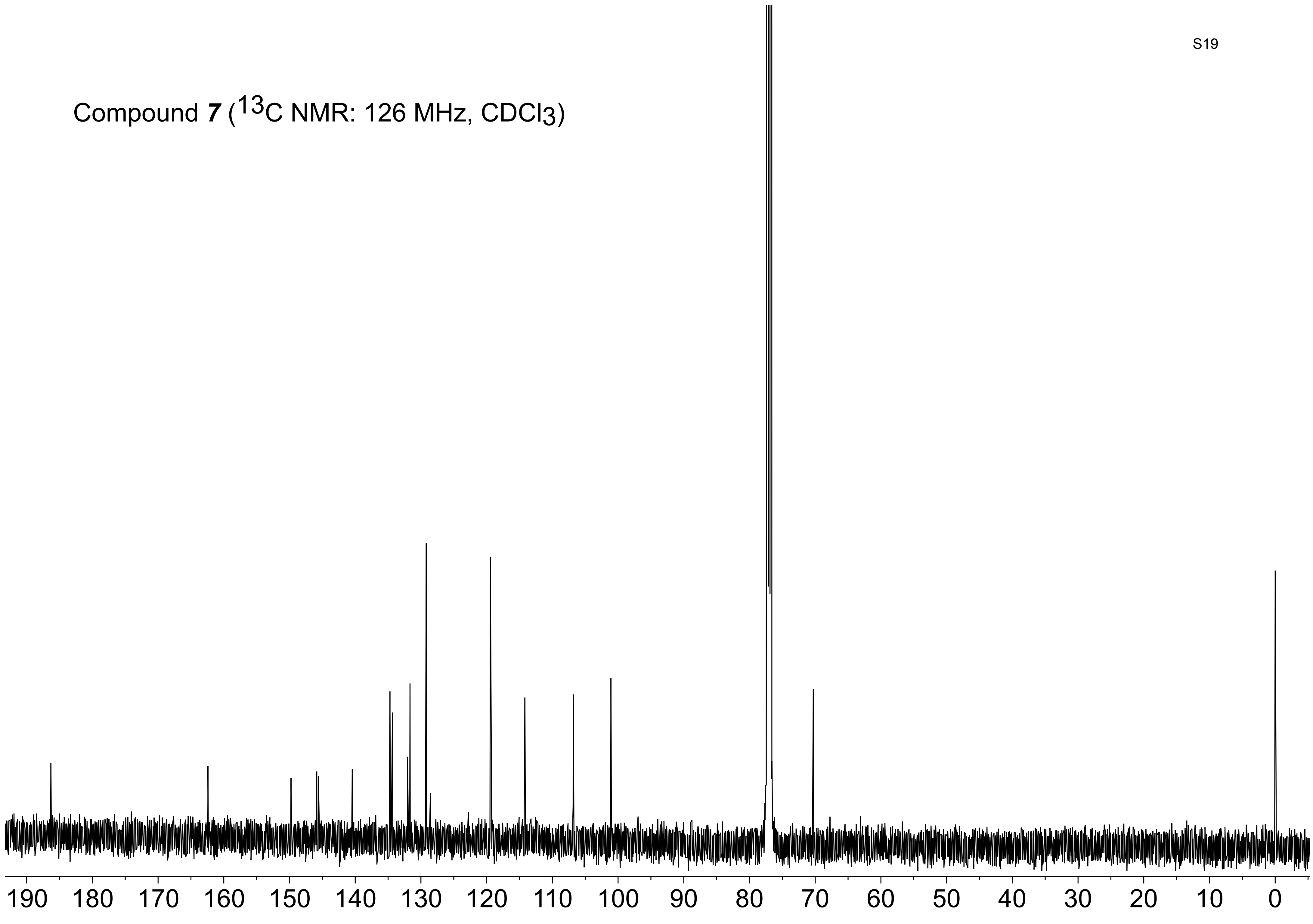

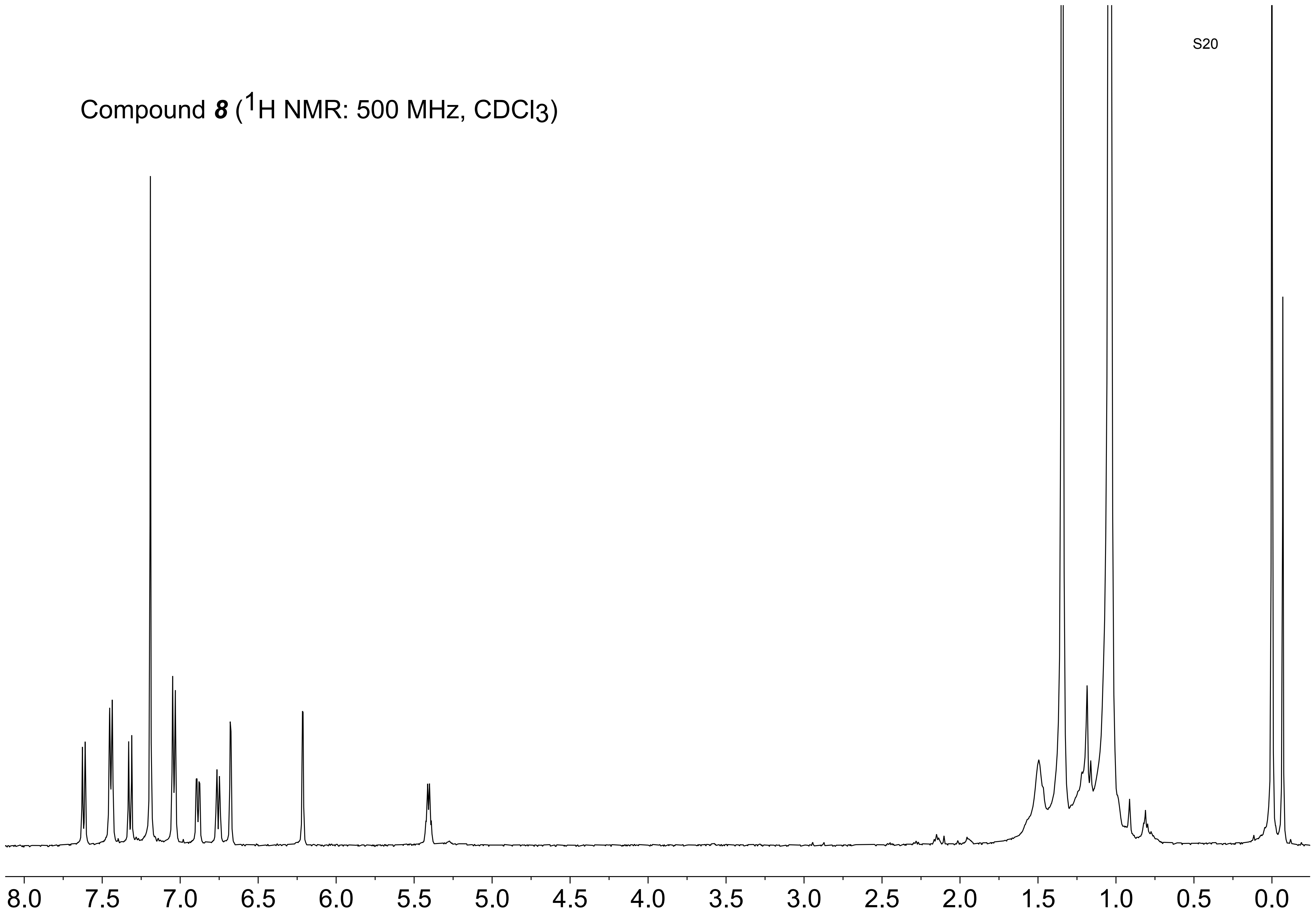
Compound 8 (1H NMR: 500 MHz, CDCl3)
8.0
7.5
7.0
6.5
6.0
5.5
5.0
4.5
4.0
3.5
3.0
2.5
2.0
1.5
1.0
0.5
0.0

Compound ***8*** ($^{19}F$ NMR: 470 MHz, $CDCl_3$)

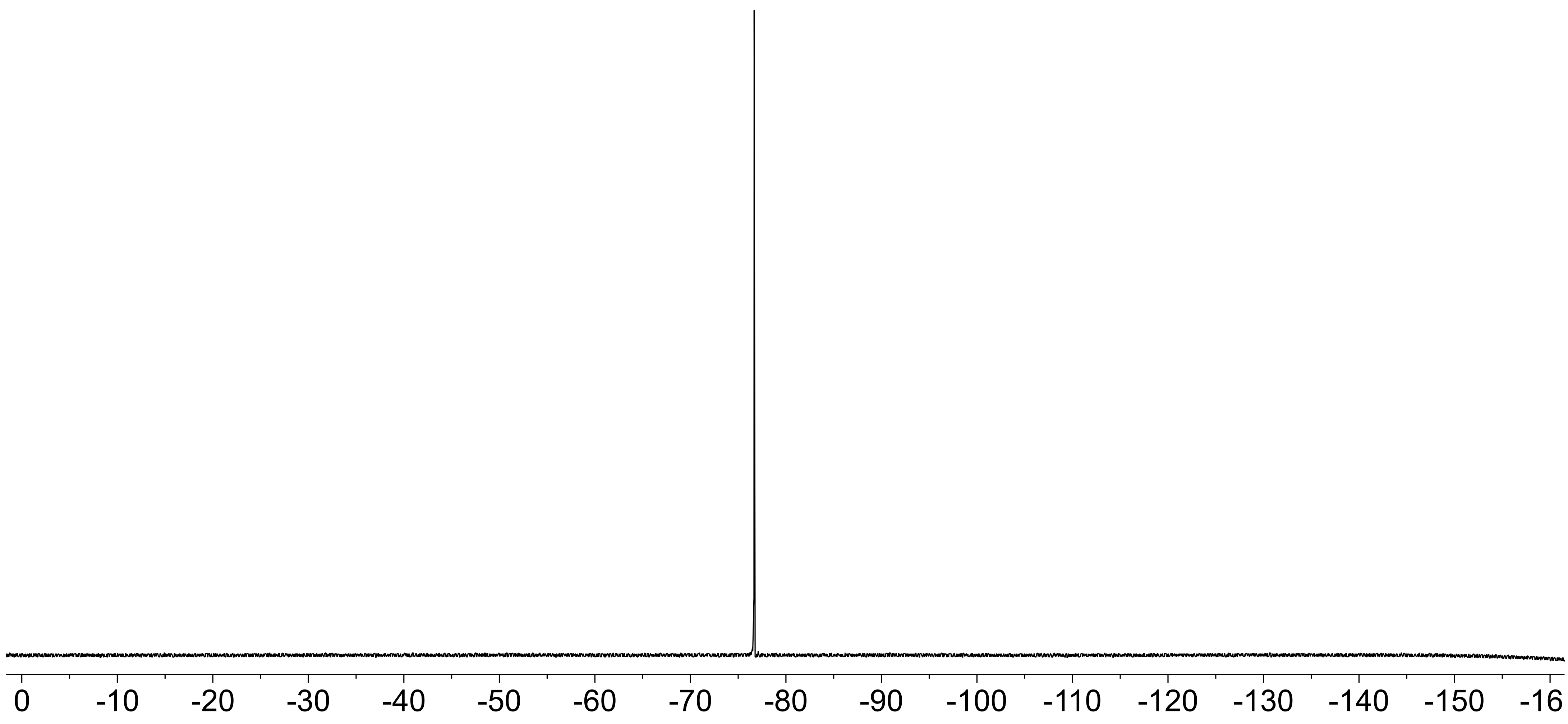

### Computational Methods and Results

As illustrated in Figure S1, the reductive debenzylation of compounds ***7*** and ***8*** may proceed through four potential reaction mechanisms. In their initial reduction reactions with $H_2S/HS^-$, compounds ***7*** and ***8*** can yield the neutral intermediates ***19*** and ***20***, characterized by the terminal -$NH_2$ groups, or the negatively charged ***11*** and ***12*** with the -$NH^-$ groups.[2] Subsequently, these neutral or anionic intermediates undergo either 1,6-elimination or hydrolysis processes, ultimately leading to the formation of the observed products. We evaluated these processes with DFT calculations using the Gaussian 16 program.[3] For each system, the electronic structures were described at the B3LYP level[4–6] with D3 dispersion correction[7] and 6-311++G(d,p) basis set. The aqueous environment was incorporated using the polarizable continuum model with a dielectric constant of 78.4.[8] The resulting reaction free energies, $\Delta G_{rxn}$, and the activation free energies, $\Delta G_a$, of these potential pathways are summarized in Table S1.

For the neutral intermediates ***19*** and ***20***, 1,6-elimination reactions exhibit $\Delta G_{rxn}$ values of 10.8 kcal/mol and 16.0 kcal/mol, respectively. The large and positive $\Delta G_{rxn}$ indicate that these reactions are unlikely to occur spontaneously at ambient conditions. Therefore, we did not search for the transition states and obtain the activation energies for 1,6-elimination reactions of neutral species ***19*** and ***20***. In comparison, $\Delta G_{rxn}$ values for the hydrolysis reactions are 2.2 kcal/mol and -2.0 kcal/mol, respectively. Despite their favorable reaction Gibbs free energies, the substantial $\Delta G_a$ values of ~20 kcal/mol render these reactions challenging to proceed at room temperature.

When considering the anionic intermediates ***11*** and ***12***, we observe a strong preference for the 1,6-elimination reactions with $\Delta G_{rxn}$ values of -20.0 kcal/mol and -19.5 kcal/mol, respectively. Furthermore, their moderate $\Delta G_a$, which are below 5 kcal/mol, allow these reactions to proceed under ambient conditions. In contrast, $\Delta G_{rxn}$ are 3.5 kcal/mol and -0.9 kcal/mol, respectively, for their hydrolysis reactions. The $\Delta G_a$ values, which are approximately 30 kcal/mol, make these reactions highly unfavorable at room temperature. Based on these results, we conclude that the most likely reaction mechanism for the reductive debenzylation of compounds ***7*** and ***8*** involves the formation of the anionic intermediates ***11*** and ***12*** from the initial reduction step, followed by 1,6-elimination reactions.

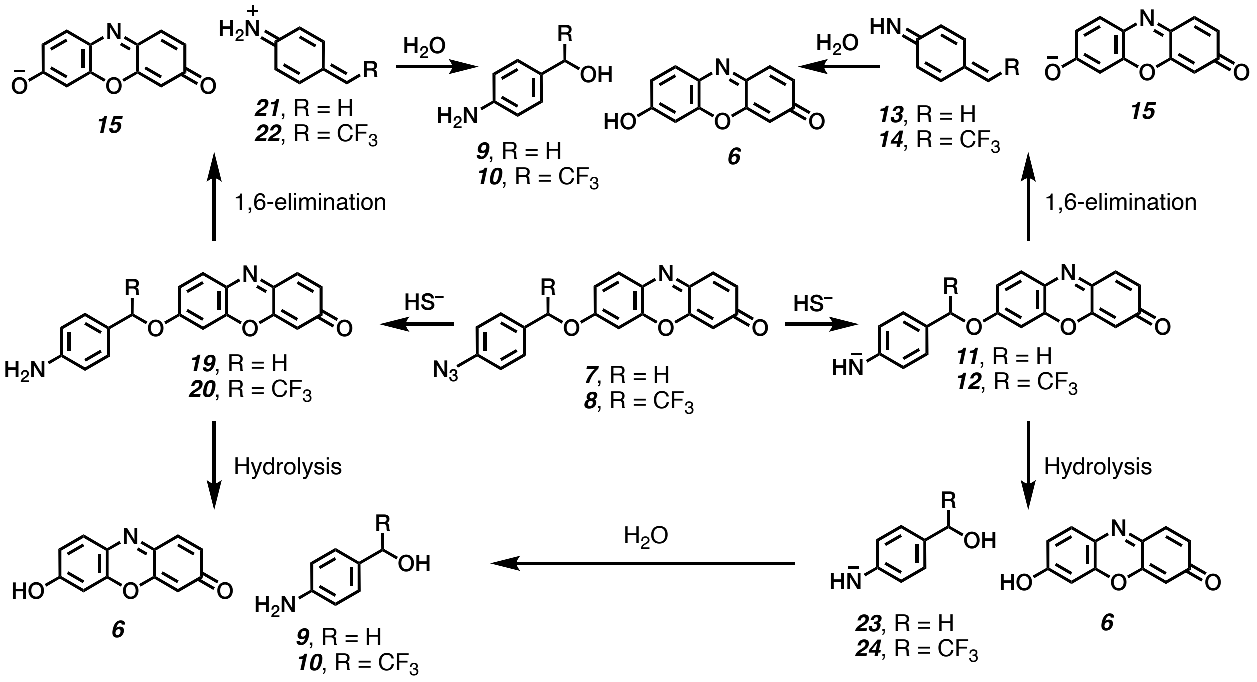

**Figure S1.** Hypothesized reaction pathways for the reductive debenzylation of ***7*** and ***8***.

**Table S1.** Reaction free energies and activation free energies for the 1,6-elimination or hydrolysis reactions of ***19***, ***20***, ***11*,** and ***12*** obtained from DFT calculations.

| Compound | $\Delta G_{rxn,\ elimination}$ (kcal/mol) | $\Delta G_{a,\ elimination}$ (kcal/mol) | $\Delta G_{rxn,\ hydrolysis}$ (kcal/mol) | $\Delta G_{a,\ elimination}$ (kcal/mol) |
|---|---|---|---|---|
| ***11*** | -20.0 | 4.4 | 3.5 | 31.7 |
| ***12*** | -19.5 | 2.7 | -0.9 | 28.7 |
| ***19*** | 10.8 | - | 2.2 | 19.3 |
| ***20*** | 16.0 | - | -2.0 | 23.7 |

### Fluorescence Measurement

**$H_2S/HS^−$ detection:** The stock solutions of probes ***7*** and ***8*** were prepared in molecular biology grade DMSO at 1 mM and 10 mM, respectively. The probes were first diluted into 10 μM concentrations with 100 mM Tris buffer (pH 7.5). Then 50 μL of the resulting probe solution (10 μM), 45 μL of Tris buffer (pH 7.5, 100 mM), and 5 μL of the reactive sulfur species were mixed in a half-area 96-well plate. Concentrations at the time of mixing were estimated to be 5 μM for the probe and 200 μM or 500 μM for the RSS. Immediately after filling, the wells were sealed with MicroAmp™ Optical Adhesive Film to avoid potential $H_2S$ gas leakage. The fluorescence of the solutions was measured (ex/em 545/585 nm) at different time intervals on a Spectra Max ID3. Note that having ≥40 nm difference in ex/em wavelengths is recommended by the instrument manufacturer).

**Limit of Detection:** Samples were prepared following the same steps mentioned above, except the $Na_2S$ solution (the $H_2S/HS^-$ source) were added at serial dilutions within a range of 0–50 μM. Fluorescence outputs were recorded after incubating for 60 minutes. Samples at each concentration were repeated for three replicates, and one-tailed Student's *t*-test was used to investigate whether the fluorescence outputs at each $H_2S/HS^-$ concentration is statistically higher than that at 0 μM (Figures 3E and 3F).

### Cell Studies

#### Preparation of Cells for microscopy imaging

HeLa cells ($3.00x10^4$) in DMEM (400 µL) with 10% FBS and penicillin (100 U/mL) were seeded into 35 mm confocal microscope dishes (VWR) and incubated at 37 °C with 5% $CO_2$. The probes (20 µM) were introduced to the cells, which were then incubated for 30 minutes. Cells were then washed with DMEM (400 µL) for 3 times to remove the probes that were not uptaken by cells, followed by the treatment of either $Na_2S$ (200 µM) or L-Cys (1 mM). For the exogenous $H_2S/HS^-$ treatments with $Na_2S$, the dishes were sealed with parafilm to form a closed-system for 5 minutes, minimizing $H_2S$ leakage. The seal was then removed and cells were incubated for the remaining hour.

#### Live Cell Imaging

CellMask Deep Red Actin dye (1 µM, 400 µL, 652/669 nm, F-actin stain) was introduced to HeLa cells in the dark following 1 hour of incubation with $H_2S$ sources. Cells were first washed with HBSS (400 µL) 3 times and the F-actin stain was introduced. Cells were then incubated for 10 minutes at 37 °C and washed out with HBSS (400 µL) 3 times after which they were used for imaging. Live HeLa cells immersed in HBSS (400 µL) were imaged on a Leica TCS SP8 confocal microscope. Signals from the probes were acquired at 571 nm excitation and 584 nm of emission.

#### MTT Assay

Cytotoxicity of the probe ***7***, ***8***, and resorufin against HeLa cells was determined using a standard MTT assay. HeLa (10,000 cells/well) were seeded in a 96-well tissue culture plate and cultured overnight to allow the attachment of the cells to the surface. The cell media was then replaced with 100 µL DMEM with 10% FBS containing varied concentrations of probes and resorufin. After 24 hours, 10 µL of the 3-(4,5-dimethylthiazol-2-yl)-2,5-diphenyl tetrazolium bromide (MTT reagent) was added to each well and the plate was incubated at 37°C for 3 hours to allow formazan crystals to form. Then, 100 µL of "detergent reagent" (from the MTT assay kit) was added to the wells and the plate was incubated in dark at rt for 2 hours to achieve full dissolution of the formazan crystals. The absorbance of each well was recorded at 570 nm on a Spectra Max ID3. The OD570 value for the sample containing cell-free media (DMEM with 10% FBS) was used for background subtraction. The OD570 values were then normalized based on the samples containing the untreated cells to obtain a percentage for cell viability. Using these values, we plotted the dose-response curves of HeLa (Figure S2), which provided $IC_{50}$ values (Table S2) through a non-linear regression fitting model.

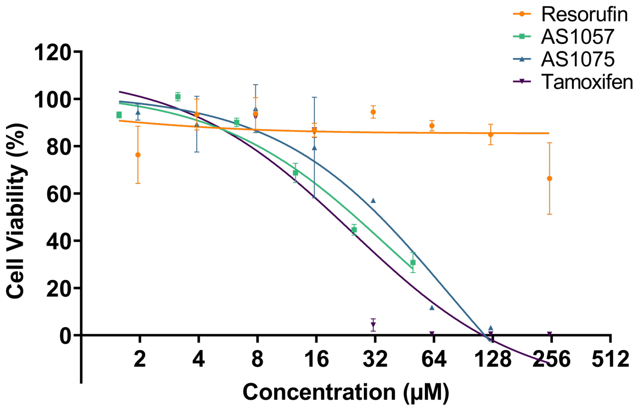


**Figure S2.** Determination of cytotoxicity.

**Table S2.** Summary of MTT results for HeLa cells

| | **Probe *7*** | **Probe *8*** | **Resorufin** | **Tamoxifen** |
|---|---|---|---|---|
| **$IC_{50}$ (μM)** | 37±23 | 76±42 | >250 | 24±23 |

### Impact of Endogenous Stimulation with L-Cys on HeLa

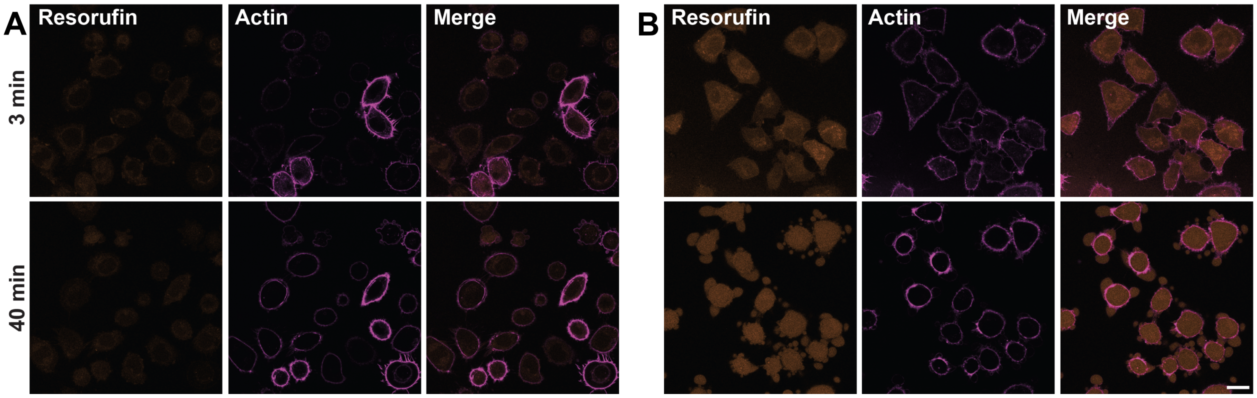


**Figure S3.** Observing cell death over 40 minutes following L-Cys treatment. Cells were incubated with either **(A)** probe **7** (20 μM) or **(B)** **8** (20 μM), and then treated with L-Cys (1 mM).

(3) Frisch, M. J.; Trucks, G. W.; Schlegel, H. B.; Scuseria, G. E.; Robb, M. A.; Cheeseman, J. R.; Scalmani, G.; Barone, V.; Petersson, G. A.; Nakatsuji, H.; Li, X.; Caricato, M.; Marenich, A. V.; Bloino, J.; Janesko, B. G.; Gomperts, R.; Mennucci, B.; Hratchian, H. P.; Ortiz, J. V.; Izmaylov, A. F.; Sonnenberg, J. L.; Williams; Ding, F.; Lipparini, F.; Egidi, F.; Goings, J.; Peng, B.; Petrone, A.; Henderson, T.; Ranasinghe, D.; Zakrzewski, V. G.; Gao, J.; Rega, N.; Zheng, G.; Liang, W.; Hada, M.; Ehara, M.; Toyota, K.; Fukuda, R.; Hasegawa, J.; Ishida, M.; Nakajima, T.; Honda, Y.; Kitao, O.; Nakai, H.; Vreven, T.; Throssell, K.; Jr., J. A. M.; Peralta, J. E.; Ogliaro, F.; Bearpark, M. J.; Heyd, J. J.; Brothers, E. N.; Kudin, K. N.; Staroverov, V. N.; Keith, T. A.; Kobayashi, R.; Normand, J.; Raghavachari, K.; Rendell, A. P.; Burant, J. C.; Iyengar, S. S.; Tomasi, J.; Cossi, M.; Millam, J. M.; Klene, M.; Adamo, C.; Cammi, R.; Ochterski, J. W.; Martin, R. L.; Morokuma, K.; Farkas, O.; Foresman, J. B.; Fox, D. J. Gaussian 16 Rev. C.01.
